# SImBA-SiQuAl: advancing high-content high-throughput phenotypic profiling of 3D microtumours

**DOI:** 10.64898/2026.04.14.718366

**Authors:** Elias Van De Vijver, Klara Dewitte, Axl Van Alboom, Christophe Ampe, Hans Van Vlierberghe, Marleen Van Troys

**Affiliations:** Department of Internal Medicine and Paediatrics; Hepatology Research Unit; Liver Research Center, Faculty Medicine and Health Care, Ghent University, Gent, 9000, Belgium; Department of Biomolecular Medicine, Faculty Medicine and Health Care, Ghent University, Ghent, 9000, Belgium; Department of Industrial Systems Engineering and Product Design, Ghent University, Ghent, 9000, Belgium; VIB-UGent Research Building FSVMII, Technologiepark-Zwijnaarde 75, Gent, 9000, Belgium

## Abstract

Three-dimensional microtumour models such as spheroids are increasingly used in cancer research as they better capture tumour architecture, growth and invasion than conventional two-dimensional cultures. However, robust and accessible tools for quantitative analysis remain limited. Here we present SImBA-SiQuAl, an integrated open-source workflow for high-throughput quantitative phenotyping of 3D spheroids and organoids. The pipeline combines SImBA, an automated image-analysis framework for performant quality-controlled image segmentation and multi-feature extraction from spheroid assays, with SiQuAl, a downstream analysis platform that automatically performs comprehensive statistical and multivariate analyses to reveal phenotypic differences between experimental conditions. In a first case study, we showcase how SImBA-SiQuAl resolves intrinsic invasion phenotypes between cancer cell lines. In a second case study, it quantifies heterogeneous responses in a spheroid drug screening assay. Together, SImBA-SiQuAl provides an effective timely tool for high-throughput, high-content microtumour phenomics in cancer research.

**MOTIVATION:** 3D-microtumour assays such as spheroids and organoids are increasingly used in preclinical research. These assays generate rich phenotypic imaging data, but automated quantitative analysis remains a major bottleneck. This limits reproducibility, scalability, and broad adoption for large-scale, high-content phenomics studies. Moreover, it impedes comprehensively addressing biologically relevant phenotypic (heterogeneous) responses in e.g. perturbation studies. SImBA-SiQuAl (**<u>S</u>**pheroid **<u>Im</u>**age **<u>B</u>**atch **<u>A</u>**nalysis - **<u>Si</u>**mba **<u>Qu</u>**antitative output **<u>A</u>**na**l**ysis) is developed to address this gap by providing an open-source, integrated workflow offering solutions in both the image processing and downstream analysis. Together, this enables in-depth quantitative analysis of 3D microtumour phenotypes across experimental settings.

## INTRODUCTION

Uncontrolled proliferation, survival, and invasive capacity are key hallmarks of cancer cells^1^ driving tumour progression and metastasis^2^. Preclinical models that quantify these functional phenotypes are essential to decipher the mechanisms of tumour progression, for drug screening or to evaluate personalized therapeutic responses. Capturing the dynamic interplay of tumour growth and invasion and the effect hereupon of drug treatment requires experimental models that extend beyond reductionist 2D systems that often overestimate drug response^4^ and cannot address phenotypes linked to multicellular 3D organisation including ECM invasion^5–14^. *In vitro/ex vivo* 3D cancer models better reflect the structural and functional complexity of solid tumours and have gained attention in preclinical research to study cancer growth, viability and metastasis^10,15–20^.

These 3D-models, here collectively termed 3D-microtumours, include multicellular tumour spheroids (MCTS) and patient-derived (tumour) organoids (PDO)^21–24^. PDOs are used to study patient-specific tumour development and progression since they offer exceptional histological and genetic fidelity, closely recapitulating the architecture and original tumour microenvironment^25–34^. Their establishment, however, remains costly, time-consuming and hard to standardise^25,35–37^, making PDOs less suitable for large scale screenings or systematic perturbation studies^38,39^. MCTS offer a complementary, experimentally more accessible 3D-model that preserves relatively complex 3D architecture, hypoxic and nutrient gradients, and multicellular organisation with in vivo-like growth kinetics and therapeutic responses^40^. MCTS can be reproducibly generated from cell lines, patient tumour tissue or heterotypic co-cultures^41–43^. Spheroids maintained in suspension serve as ideal tool for growth and survival monitoring^44^. Equally, spheroids embedded in ECM hydrogels^45–48^ serve as physiologically relevant test model to capture invasion modes and tumour-ECM interactions, providing insight in a dynamic “invasive growth phenotype”. This creates a range of opportunities for large scale perturbation analyses including gene knock-down, knockout^49–51^ or drug screenings at high throughput^52^ as reflected by the marked increase in spheroid cancer research in recent years.

Research using MCTS often relies on imaging-based phenotypic studies using different microscopic technologies including high-throughput commercial system such as Incucyte^53^, Operetta CLS^54^, CellVoyager^55^ or ImageXpress micro^56^.The MISpheroID database^57^ is a recent initiative that addresses experimental standardization for spheroid research revealing substantial heterogeneity across studies which led to the definition and promotion of minimal reporting requirements. The recent establishment of SLiMIA^58^ as the first, large-scale, open-access database dedicated to spheroid images, further emphasizes the growing interest into and the future potential of spheroid research. Although these efforts are improving experimental consistency, the greater bottleneck of spheroid/organoid research lies in the downstream image processing and quantification strategies. Current methods rarely exploit the wealth of information that 3D microtumour assays inherently provide. Remarkably, many studies still rely on manual or semi-automated segmentation, a restrictive, labour-intensive approach that often limits image-based output to a set of rudimentary features such as area, perimeter, or radius^50,59–65^. The field thus still lacks accessible, reproducible pipelines that assist users to easily transform raw image data into high quality, high-content phenotypic data. This lack reduces scalability, impedes reproducibility, and more essentially, prevents the discovery of subtle but biologically relevant differences between tested conditions. Critically, this also prevents systematic integration of detailed functional phenotype data (phenomics) with increasingly available cancer multi-omics datasets and systematic contributions to understanding the determining role of phenotypic heterogeneity and clonal divergence in cancer treatment resistance and metastasis^66–68^.

To avoid that the majority of information present in 3D-microtumour-based studies remains underexplored, user-friendly, integrated pipelines that bridge the gap between complex image datasets and high-content, multiparametric readouts need to be established^24^. Several reported tools have addressed specific aspects of this challenge including AnaSP ^69^, SpheroidSizer^70^, SpheroidJ^71^, INSIDIA(2.0)^72,73^, TASI^74^, SpheroScan^75^, SperoidAnalyzeR^76^, ReViMS^77^ and ReViPS^78^. The features, capabilities and limitations of these available tools are listed in Table 1. Whereas most of these provide useful functionalities for spheroid segmentation and/or basic feature extraction, they typically focus on a narrow set of metrics, require manual pre-processing or already segmented input, are optimized for specific cell types or imaging conditions or lack end-to-end automation. Importantly, none of them provide integrated statistical and multiparametric analyses. Consequently, the phenotypic richness of microtumour models still remains insufficiently exploited.

**Table 1.**
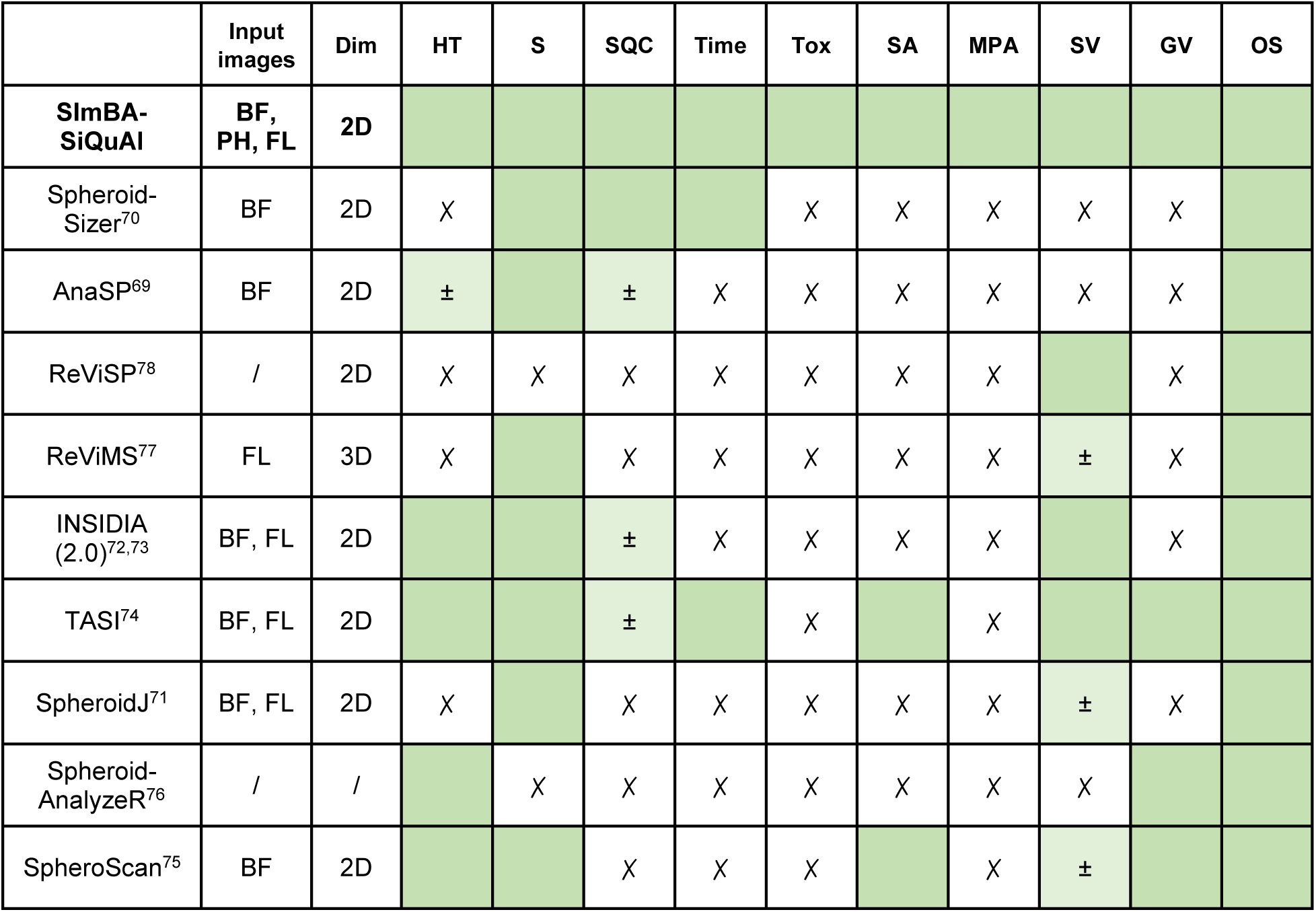
Comparison of available open-source software tools to analyse spheroid assays. BF: Brightfield; PH: Phase-contrast; FL: Fluorescent; Dim: Input image dimension; HT: High throughput; S: Segmentation; SQC: Segmentation Quality Control; Time: Time-series analysis; Tox: cytotoxicity assessment; SA: Statistical analysis; MPA: multiparametric analysis; SV: Segmentation visualisations; GV: Graphical Visualisations; OS: Open-source.

We introduce SImBA-SiQuAl, an open-source, automated, end-to-end analysis workflow for robust quantification of times series of single 3D spheroids or organoids. SImBA (<u>S</u>pheroid <u>Im</u>age <u>B</u>atch <u>A</u>nalysis) performs accurate, automated segmentation across bright-field, phase-contrast and fluorescent spheroid images and extracts an extensive set of metrics for growth (kinetics), invasion, morphological complexity and viability. Using this numeric output of SImBA, SiQuAl (<u>SI</u>mBA <u>Qu</u>antitative output <u>A</u>nalysis), seamlessly implements univariate statistical testing, multivariate profiling and clustering, combined with graphical output, enabling data-driven comparisons of phenotypic states linked to treatment responses with clear biological interpretability.

In this report, we present the software features, segmentation performance, varied output and data analysis strategies. Two case studies are used to showcase the wealth of data produced by the software tandem and illustrate its capacity to uncover nuanced treatment effects.

## RESULTS

### The SImBA-SiQuAl workflow enables end-to-end quantitative spheroid phenotyping

The SImBA-SiQuAl tandem presented here is developed to systematically transform raw spheroid image time series from different experimental conditions, into high-dimensional, quantitative, phenotypic descriptions that allow biological interpretation. SImBA-SiQuAl is designed for image-based datasets aimed at comparing spheroid growth and/or invasion over time in response to perturbations such as drug treatment, growth factor stimulation, gene knockdown or knockout, or protein overexpression, at any level of throughput. SImBA performs image-processing on spheroids acquired in time, includes segmentation quality-control (QC), has no limitations in number, nature or source of images, generates visual output and quantifies multiple features. SiQuAl subsequently processes the quantitative output for downstream statistical comparisons and graphical visualization.

To ensure broad adoption, both tools are open source. SImBA is developed as a macro in FIJI^79^, a widely used microscopy image analysis platform, whereas SiQuAl is a Python-based pipeline that can also be run as a standalone executable for users without programming experience.

Here, we first provide an overview of the SImBA-SiQuAl workflow (Figure 1), highlighting key analytical steps. Note that a comprehensive description of the SImBA-SiQuAl workflow is detailed in Supplementary Materials. Additionally, the SiQuAl analysis of both case studies reported below will be available together with two guided examples and a user manual and for first-time users^80,81^. Starting from a standardized folder organization (Figure 1: step 1, Figure S1A), users first define the biological context of the experiment. SImBA offers either a growth or invasion analysis (step 3A-B). *Growth* analysis (A) is suited for spheroids cultured in medium whereas *Invasion* analysis (B) is intended for spheroids embedded in a 3D-hydrogel. This choice determines whether segmentable objects outside the main spheroid mass are considered biologically relevant or not, thereby ensuring their removal in *Growth* analyses and inclusion in *Invasion* analyses (detailed in Supplementary Materials 1.1 and Figure S1B). This unique flexibility substantially expands SImBA’s applicability across diverse microtumour studies.

**Figure 1.**
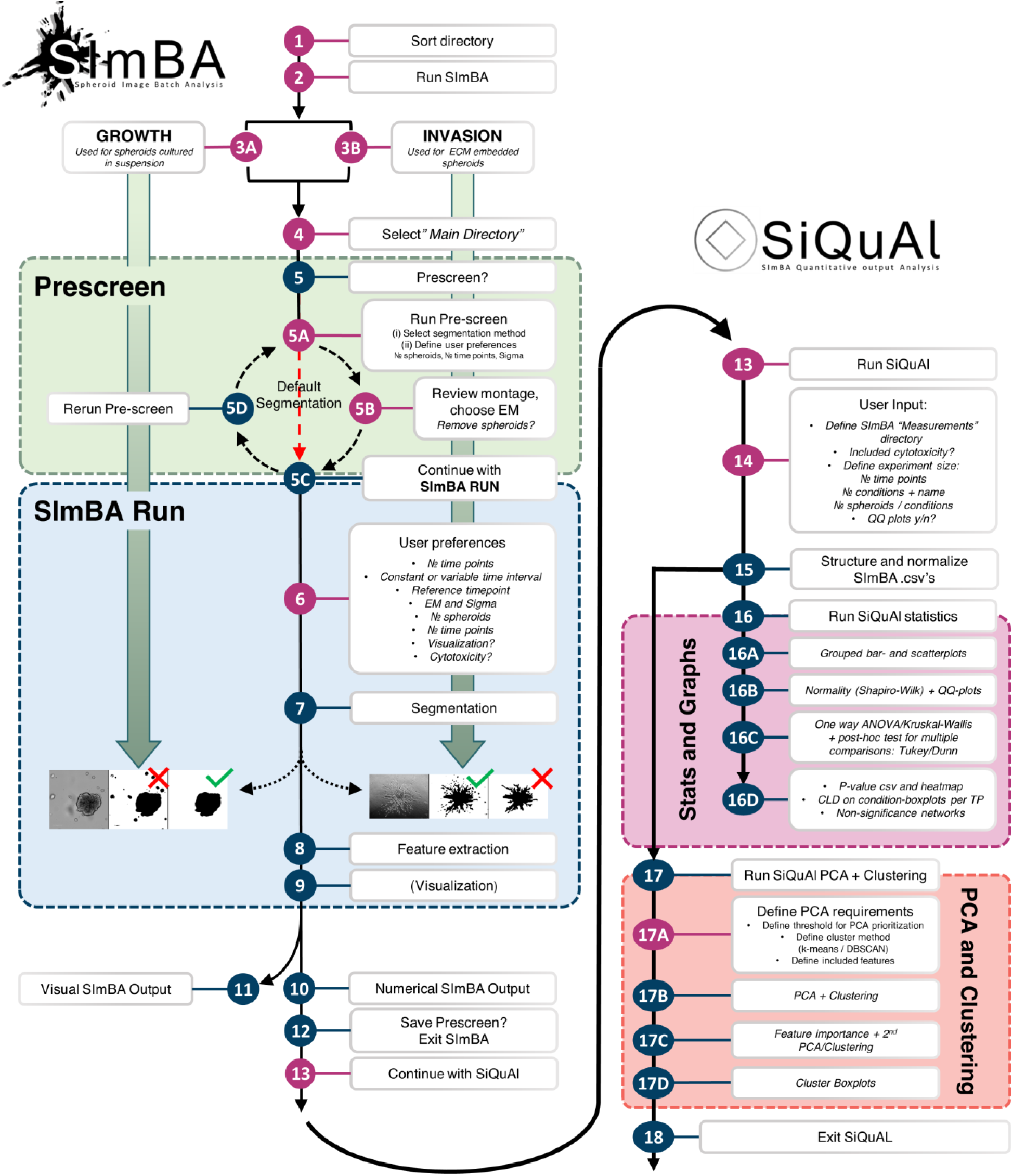
End-to-end SImBA-SiQuAl workflow for high-content 3D microtumour phenotyping. Spheroid image time sequences of both growth and invasion assays at any throughput are processed by **SImBA** including a unique segmentation quality-control step (Prescreen), generating visual outputs and quantitative data for 29 spheroid features (see Figure 2A and Table S1). The numerical output is analysed by **SiQuAl** through both single-parameter statistical testing and multivariate analysis using PCA and cluster analysis. EM: error margin; Sigma: gaussian smoothing control, Pink nodes: user input/choice, blue nodes: fully automated.

A uniquely distinguishing central attribute of the SImBA workflow is the *Prescreen* module (step 5) which enables essential QC prior to full-scale SImBA segmentation and quantification. During this QC-step, users can iteratively compare five distinct segmentation methods, using easily surveyable segmentation montages (detailed in Supplementary Materials 1.1).

The *Prescreen* also enables fine-tuning of additional parameters such as level of Gaussian smoothing by adjusting the Sigma-value. Concentric reference rings in the *Prescreen* output assist users in defining an image-specific error margin (EM) (Figure S1C). This EM provides an additional safeguard for high-quality segmentation since it allows to exclude artefacts, debris or neighbouring spheroids present in the periphery of images that would skew the analysis. Thereby, the EM allows more images to be retained for downstream analysis. An in-depth description of *Prescreen* functionalities is provided in Supplementary Materials 1.1 and Figure S1C. While the *Prescreen* increases run-time, omitting this step would necessitate manual image-by-image QC, individual segmentation assessment or separate image pre-processing, which are considerably more labour-intensive, time-consuming, less standardised or scalable and which can introduce unintended bias. By enabling segmentation strategies and parameters to be tested across the full dataset, the *Prescreen* module greatly improves consistency, reproducibility, and confidence in downstream quantification beyond what is feasible by prior manual inspection. This explicit QC layer, rarely incorporated in existing spheroid analysis software, is a key strength of SImBA.

Following *Prescreen*, segmentation settings are defined and, after experimental annotation (Figure 1, step 6), SImBA proceeds fully autonomously to batch-segment all spheroid time-series (7), to extract a multidimensional quantitative feature set (8) and to generate visuals (9). In total, for each spheroid numerical values are extracted for 29 features in five categories, as shown in Figure 2A. A first feature category captures spheroid size metrics over time, either including or excluding invading areas, as well as features that quantify spheroid growth kinetics. Second, spheroid invasion is described through multiple complementary features. Additionally, measures of spheroid architecture complexity (over time) in the form of different area/perimeter ratios, measures of spheroid overall morphology and (treatment-induced) cytotoxicity are also extracted. Table S1 lists the mathematical definitions of the extracted quantitative features and their biological relevance. Next to this quantification, six types of visual output are provided for each spheroid (see Figure S1D, Table S2).

**Figure 2.**
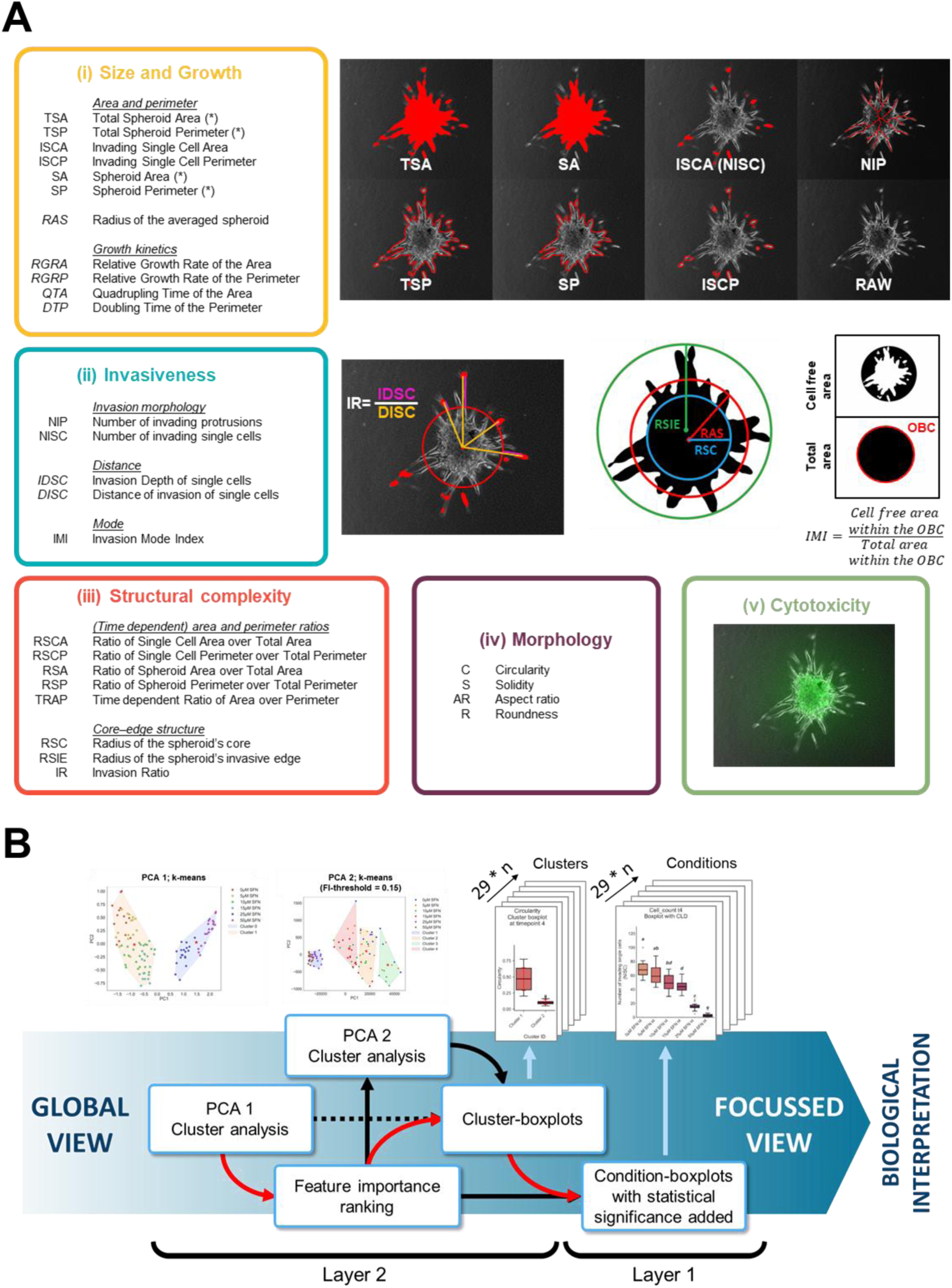
SImBA-SiQuAl feature extraction and 2-layered phenotypic analysis workflow. (A) SImBA feature categories (i) Size and Growth: 11 metrics of areas, perimeters and kinetic changes; (*) after normalization (N) to t_0_ reference by SiQuAl indicated by e.g. NTSA: normalized total spheroid Area, (ii) Invasiveness: 5 features describing invasion morphology, single-cell distance metrics and invasion mode, (iii) Structural Complexity: 8 features describing (time-dependent) spheroid architecture. (iv) Morphology: 4 global shape descriptors, (v) 1 feature of cytotoxicity: mean intensity of toxicity staining in the TSA. For complete feature descriptions: Table S1. Numerical values are reported in eight csv files in the SImBA Measurement folder (Table S2). (B) PCA1 and clustering of the full feature set provide an unbiased multivariate overview. SiQuAl ranks feature importance and performs PCA2 with clustering excluding low-impact features. Cluster-boxplots and condition-boxplots for all features are used together to identify which features drive phenotypic separation and support biological interpretation as indicated by the arrows (n = timepoints).

Importantly, whereas conventional pipelines typically stop after segmentation or feature extraction, SImBA seamlessly extends into SiQuAl for comprehensive quantitative analysis and associated statistics (Figure 1: step 13-18). After input of experimental meta-data (14), SiQuAl operates fully independent. It first reorganises the numerical feature data and, where appropriate, normalises values to the starting reference spheroid at t_0_ for each time series (15).

SiQuAl then performs two complementary layers of analysis (step 16-17) (see Figure 2B). First, it conducts a comprehensive statistical evaluation of all extracted features at each time point (16). This includes normality testing (16B), which determines the choice of parametric or non-parametric statistical comparisons across conditions (16C). Layer 1 results are visualised per feature in several ways (16D): condition-boxplots per timepoint including statistical outcome shown as compact letter display (Figure 2B), p-value matrices with corresponding annotated heatmaps and non-significance networks summarising relationships between experimental conditions (Supplementary Materials 1.2, Table S3). Together, this already provides a detailed overview of how individual spheroid features change across experimental conditions over time.

Second, and unique within spheroid image analysis software, SiQuAl integrates by default all features per time point into a single multivariate analysis (step 17, Figure 2B layer 2). By performing principal component analysis (PCA), SiQuAl enables direct comparison of spheroids based on combined growth, invasion, complexity, morphology and viability characteristics. Subsequent clustering (either k-means or DBSCAN), enables the identification of distinct spheroid subsets that differ significantly across the full parameter set (17B). Importantly, SiQuAl not only groups spheroids into phenotypic clusters but also allows easy assessment of why clusters form. It does this by ranking features contributing most to the clustering by means of a feature-importance (FI) value attributed to each feature (17C). FI is the weighted mean of all absolute principal component loadings weighted by their explained variance (detailed in Supplementary Materials 2). Moreover, SiQuAl also runs a secondary PCA (17C) where only features with an FI higher than a set threshold are retained. In doing so, features of lesser importance, with lowest clustering impact, are removed to improve the group separation. To aid biological interpretation of the cluster output, SiQuAl produces, for both the initial and secondary PCA, cluster-boxplots (17D) that visualise how (key) features differ between clusters at each timepoint. This enables rapid global identification of the features driving cluster separation. This allows focus on the condition-boxplots (from the first analysis layer) for these features to assess the underlying statistical differences between experimental conditions. The added value of these two tuneable and complementary layers of the SiQuAl analysis (global and focussed) is depicted in Figure 2B. By explicitly linking clusters to key features on the one hand and to statistical differences between conditions in these features on the other hand, each cluster is translated into a specific functional phenotype. This forms a robust basis for detailed biological interpretations. How the multi-layered high-content output can be optimally, interactively exploited is further illustrated in the results walkthrough in Supplementary Materials 2 and in the case studies below.

Coupling SImBA to SiQuAl thus provides a single workflow from automated image segmentation to data interpretation allowing both univariate statistical- and multiparametric, phenotypic analysis over time with manuscript-ready visualisations. Note that the SiQuAl analysis is fast (e.g. 7 minutes for case study 2 below) and can easily be run consecutively using a different clustering strategy or FI threshold. A complete overview of all output produced by SImBA-SiQuAl and their structure within the main directory is provided in Tables S2 and S3.

Below we demonstrate the performance, analytical depth and application potential of SImBA-SiQuAl. First, the robustness of SImBA segmentation is benchmarked against manual expert annotations. Subsequently, we present two representative application scenarios (case study 1 and 2). These illustrate the actual output that SImBA-SiQuAl generates and its capacity to support relevant phenomics-based interpretations.

### SImBA segmentation is robust across imaging modalities and matches expert annotation

To substantiate SImBA’s segmentation performance we evaluated ∼3500 spheroid images including phase-contrast, brightfield and fluorescence images (Figure 3A). The images are either obtained from the public SLiMIA database^82^ or newly generated (for case studies below). SImBA segmentation performance is summarized in the stacked bar plot of % segmentation accuracy (Figure 3B). See Table S4 for a detailed overview of image acquisition settings, SImBA segmentation parameters and performance. Prescreen-based SImBA segmentation QC shows high accuracy across all modalities: 2653/2712 phase-contrast images could be segmented accurately (97.82%), 605/613 brightfield images (98.69%), and 170/171 fluorescence images (99.42%), yielding an overall accuracy of 98.05%.

**Figure 3.**
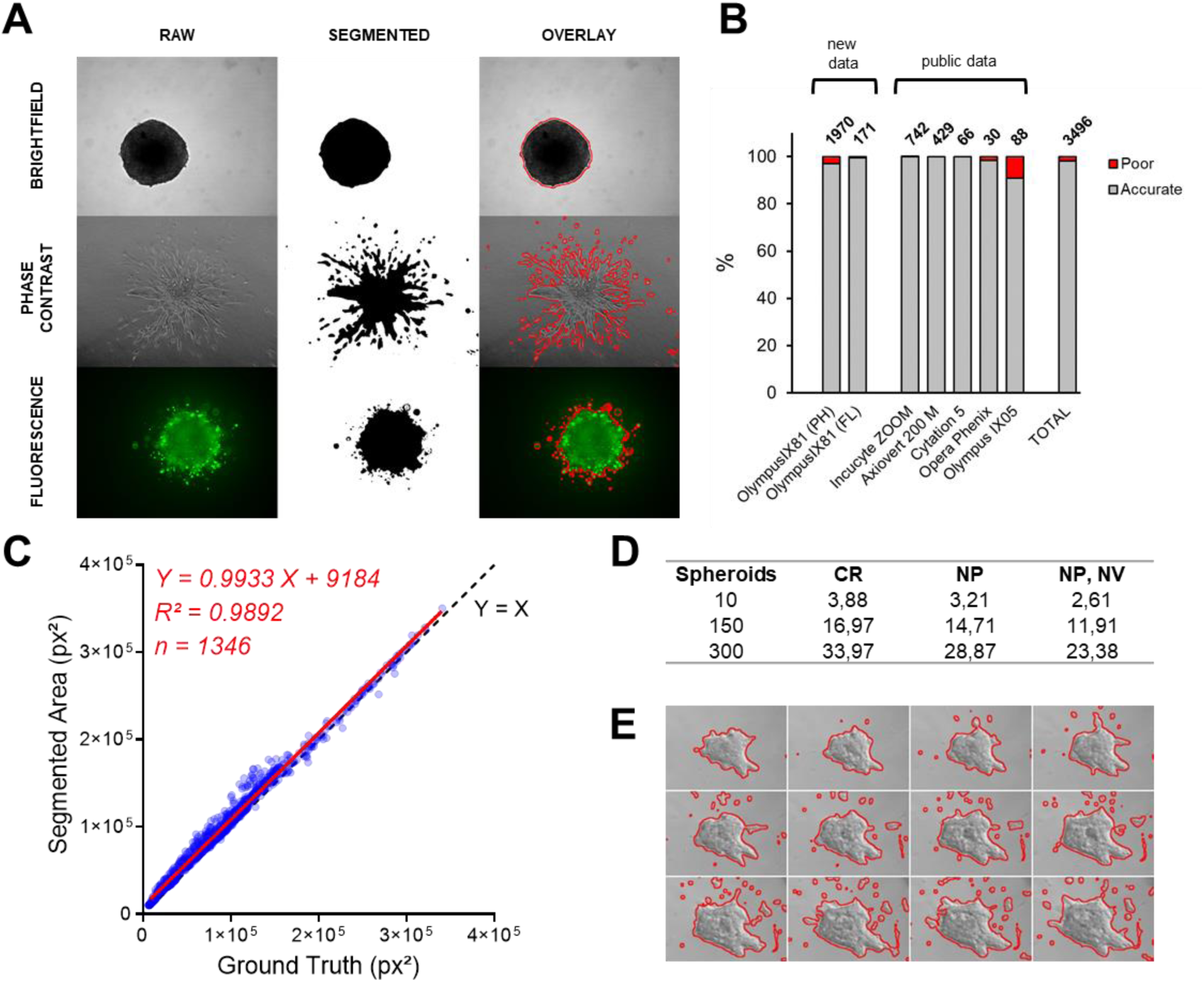
SImBA segmentation performance across imaging modalities and datasets. (A) Representative SImBA segmentation on brightfield, phase-contrast and fluorescence images. Left to right: Raw image, binary mask, overlay with SImBA-segmentation (red contour). (B) Stacked bar plot summarizing the percentage of accurately (grey) and poorly segmented (red) on images from the public SLiMIA database (per imaging system) or generated in this study (Table S4). (C) Area (px²) measured by SImBA versus manually derived ground truth for 1,346 spheroids (SLiMIA) red line: linear regression fit; black, dashed line: Y = X). (D) SImBA runtimes in minutes; CR: complete run, NP: no Prescreen, NP, NV: no Prescreen, no visualisation. (E) Example of SImBA segmentation (red outlines) across a time series of an ECM-embedded invasive murine mammary organoid (Cell Image Library RRID:SCR_003510).

To further benchmark SImBA’s automated segmentation, we compared SImBA segmentation of 1346 images of the public set to their manually segmented *ground truth* masks, available in SLiMIA^82^ (Figure 3C,S2). Area measurements showed near-perfect agreement with manual annotations, with a slope not significantly different from 1 (p = 0.7369) (Figure 3C). Similarly, perimeter measurements also strongly correlated (Figure S2B). These data indicate that automated segmentation of spheroid images matches expert manual annotation and can thus be confidently used for unbiased quantification. Moreover, SImBA segmentation and feature extraction is fast as documented in Figure 3D. Finally, we used SImBA to successfully segment ECM-embedded organoids (Figure 3E and S2C-D), supporting that SImBA can be applied to both spheroid and organoid models without adaptations.

### Case study 1: SImBA-SiQuAl identifies distinct spheroid invasion phenotypes

In a first case study, we assessed the ability of SImBA-SiQuAl to distinguish distinct invasion phenotypes. The case study contains spheroids of eight cancer cell lines, selected to ensure a diverse range of intrinsic invasive behaviours (cell lines and experiment size details: Table S5). Spheroids were embedded in collagen type I 3D-ECM and monitored over three consecutive time points with 24h intervals (t_0_, t_1_, t_2_) to quantify phenotypic differences over time.

SImBA (using default segmentation, Sigma at 6.5) and SiQuAl results are shown in Figures 4 and S3. In line with the strategy in Figure 2B, we first assessed the multiparametric PCA-clustering (Figure 4A-B) to capture the global phenotypic variation within the dataset. Using feature-importance (FI) ranking (Figure 4C), this is focused to feature-level, by consecutively using the cluster- and condition-boxplots (Figure 4D-E) at each time point to pinpoint which features drive phenotypic separation.

**Figure 4.**
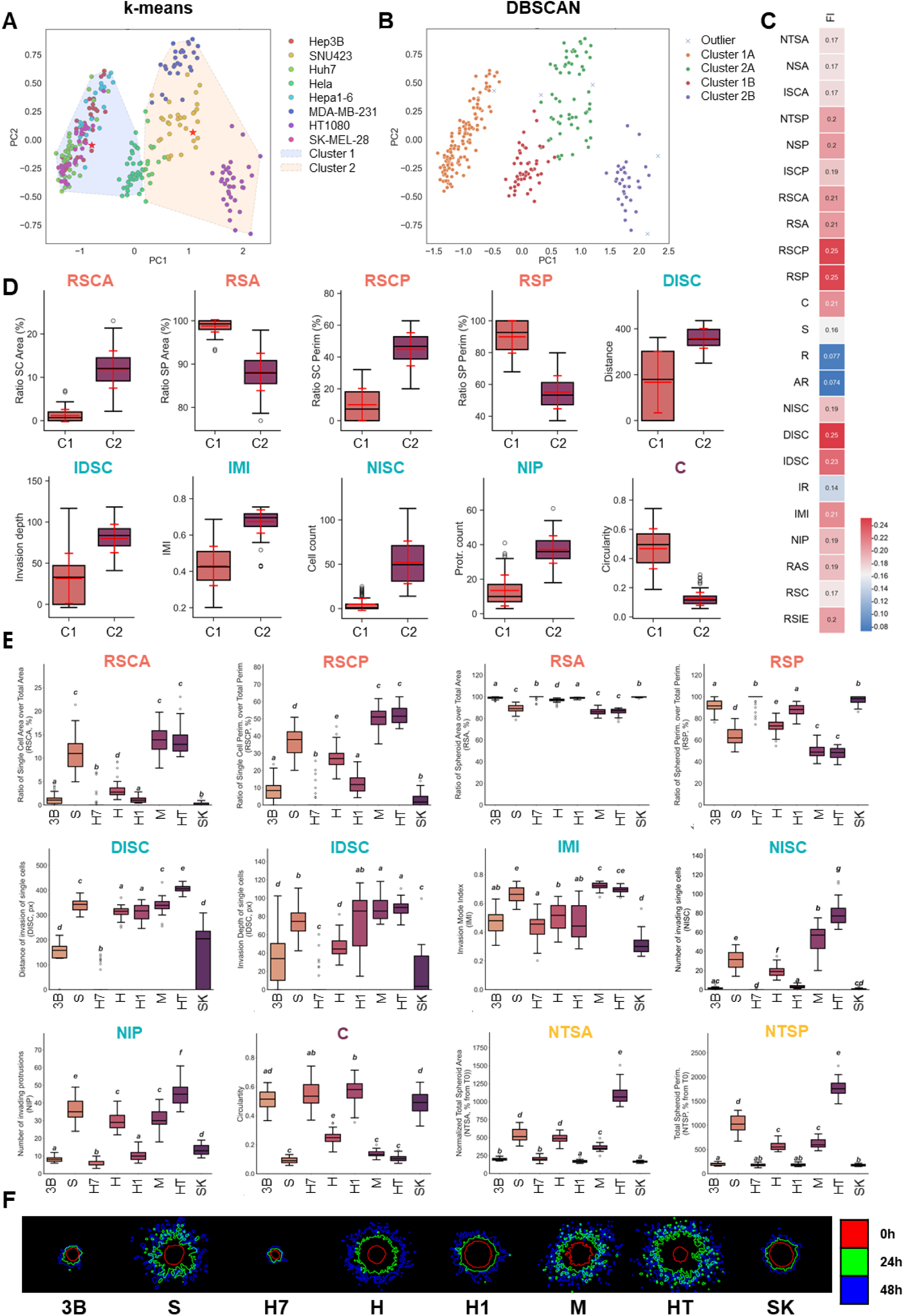
Case study 1: SImBA-SiQuAl resolves distinct spheroid invasion phenotypes. (A) PCA1 using all features with k-means clustering after 24h (t_1_). Convex hulls indicate k-means cluster boundaries, stars indicate cluster centres. (B) DBSCAN clustering at t_1_ identifies four phenotypic clusters; crosses indicate DBSCAN-identified outliers (n=6). (A-B) points represent individual spheroids. (C) FI of PCA1 at t_1_ (blue: low, red: high). (D) k-means cluster-boxplots of key FI-ranked features: ratios of single-cell/spheroid to total area and perimeter (RSCA, RSCP, RSA and RSP), single cell distance of invasion (DISC) and invasion depth (IDSC) and invasion mode index (IMI), number of invading single cells (NISC), protrusions (NIP) and circularity (C) at 24h; C1: Cluster 1, C2: Cluster 2; Mean with SD in red. (E) Condition-boxplots of key FI-ranked features at 24h: RSCA, RSCP, RSA, RSP, DISC, IDSC, IMI, NISC, NIP, C, and normalised total spheroid area and perimeter (NTSA, NTSP). Compact letter displays indicate statistically non-significant groups (ANOVA + Tukey or Kruskal–Wallis + Dunn /BYK based on normality). (D-E) Boxplots indicate median and IQR with outliers. (F) Representative SImBA segmentation overlays showing spheroid outlines over time using colour code. (E-F) 3B: Hep3B; S: SNU423; H7: Huh7; H: Hela; H1: Hepa1-6; M: MDA-MB-231; HT: HT1080; SK: SK-MEL-28.

K-means clustering on PCA1 identified two major clusters after 24h (Figure 4A). *Cluster 1* comprised Hep3B, Huh7, Hepa1-6, and SK-Mel-28 spheroids and *Cluster 2* SNU423, HT1080, MDA-MB-231. Hela spheroids, central in the PCA, clustered with *Cluster 1*. To explain the observed clustering patterns, we examined features driving phenotypic separation through FI (Figure 4C). PCA-clustering was primarily driven by structural complexity and invasiveness features including amongst others *single-cell/spheroid to total area and perimeter ratios* (RSCA, RSCP, RSA and RSP), *distance of invasion (DISC)*, *invasion depth of single cells* (IDSC) and *invasion mode index* (IMI). K-means cluster-boxplots (Figure 4D) revealed markedly higher values in *Cluster 2* for these top-ranked features, including greater DISC, IDSC, RSCA, RSCP, and increased IMI, indicating that for this cluster single cell invasion is more prominent, and single cells invade the 3D matrix farther. Concurrently, *Cluster 2* is also characterised by an increased number of invading single cells (NISC) and protrusions (NIP), combined with lower circularity (C) (Figure 4C-D). Combined, these results clearly define *Cluster 2* as exhibiting a more invasive phenotype after 24h, whereas k-means *Cluster 1* comprised low-invasive spheroids.

Complementary DBSCAN clustering on PCA1 (Figure 4B) further subdivided the two main clusters obtained by k-means, into four density-based clusters (for clustering method differences: Supplementary Materials 1.2). Low-invasive k-means *Cluster 1* was divided into 2 clusters here annotated as *Cluster 1A* and *1B*, while the invasive *Cluster 2* separated into *Cluster 2A* and *2B*. DBSCAN cluster-boxplots (Figure S3A) demonstrated that *Cluster 1A* corresponds to a compact, non-invasive phenotype, with very low DISC and IDSC, near-maximal RSA and RSP, negligible NISC and NIP, and low IMI values. *Cluster 1B* displayed intermediate values across these metrics and thus represents a low-invasive phenotype, consistent with its overlap with the low-invasive k-means *Cluster 1*. In contrast, *Cluster 2A* and *2B* both reflect invasive phenotypes and display marked increases in invasion-associated parameters. *Cluster 2B* is further distinguished from *2A* by substantially larger total spheroid area and perimeter (*NTSA, NTSP*), associated growth-related metrics (*not shown)*, and the highest NISC, DISC and NIP (Figure S3A), indicating the most aggressive invasion phenotype within the dataset.

This clearly demonstrates the added value of the flexible SiQuAl set up, including the use of different clustering modalities. Indeed, k-means clustering here provides a broad separation into low- and a high-invasive capacity, while DBSCAN refines this further into four biologically distinct phenotypes: a compact, non-invasive subtype (*Cluster 1A*), a protrusion-dominated but relatively circular low-invasive subtype (*Cluster 1B*), an invasive subtype (*Cluster 2A)*, and a highly invasive subtype characterised by extensive growth and the highest degree of single-cell invasion (*Cluster 2B*).

SiQuAl also provides condition-boxplots that, in this case, ‘deep-translate’ the cluster-associated features into cell line-specific phenotypic profiles. Global analysis identified Hep3B, Huh7, Hepa1-6 and SK-Mel-28 as predominantly non-invasive cell lines (Cluster 1, 1A; Figure 4A-B), characterized by an overall significantly lower invasiveness, with higher circularity and low NISC. The increased distances (DISC, IDSC) observed for Hepa1-6 here only reflects invasion by a small number of cells rather than a global invasive phenotype (Figure 4E). In contrast, SNU-423, MDA-MB-231 (Cluster 2A) and HT-1080 (Cluster 2B) displayed significantly increased invasion-associated features, confirming their invasive phenotype. The condition-boxplots further allowed invasion-associated features to be interpreted at the condition level, by showing that HT-1080 had higher NISC, DISC, NIP, NTSA and NTSP values than SNU-423 and MDA-MB-231, indicating a more pronounced invasive profile (Figure 4E). HeLa spheroids displayed an intermediate phenotype with significantly higher invasion than the non-invasive cell lines but considerably lower values than the highly invasive cell lines. These phenotypic similarities and differences on condition level are further supported by the parallel-coordinate plots in Figure S3B which provides a scaled overview of all features across all spheroids. This clearly visualizes the subdivision of low-, intermediate and (highly) invasive phenotypes/cell lines.

After 48h of invasion (Figure S3C-D), clustering resolution decreased as continued spheroid growth and matrix invasion (blue outline, Figure 4F) led to phenotypic convergence among cell lines, causing DBSCAN to resolve only two major clusters (high- vs low-invasive) similar to k-means clustering. Note however that the secondary PCA restricted to high-impact features partially restores phenotypic separation (Figure S3E), again underscoring the analytical resolving power of SImBA-SiQuAl. In addition, this also highlights the need for multiple timepoints.

Together, these results demonstrate that SImBA-SiQuAl differentiates distinct spheroid invasion phenotypes through multiparametric clustering while providing a flexible and transparent framework to interpret the specific features driving phenotypic divergence over time. Although an analysis limited to individual parameters such as total spheroid area or perimeter (NTSA, NTSP) (Figure 4E) already reveals distinct invasive capacity of the tested cells lines, these individual parameters cannot recapitulate the same distinctions revealed by the integrated multiparametric analysis and underlines that detailed discovery-oriented phenomic interpretation cannot rely on only one or a few features. This case uses distinct cell lines, but the accurate resolution of detailed invasion phenotypes by the SImBA-SiQuAl framework highlights the potential to quantify phenotypic heterogeneity within complex data-sets from high-throughput screens or from heterogeneous clonal tumour populations (see discussion).

### Case study 2: Capturing Distinct Sorafenib-Induced Phenotypic Responses in HCC Cell Lines

As second case study, we used SImBA-SiQuAl to evaluate treatment-induced phenotypic changes in spheroids of two hepatocellular carcinoma (HCC) cell lines (SNU423, Hep3B). The cytotoxic, tumour growth-inhibiting drug sorafenib (SFN), standard of care for advanced HCC for decades^77^ was used to treat matrix-embedded spheroids with a range of concentrations (0-50µM SFN), which were imaged (phase contrast) every 24h from 0 to 96h. SFN-induced cytotoxicity was assessed at 96h by SYTOX-Green staining. Results are presented in Figure 5 and 6; metadata in Table S5.

**Figure 5.**
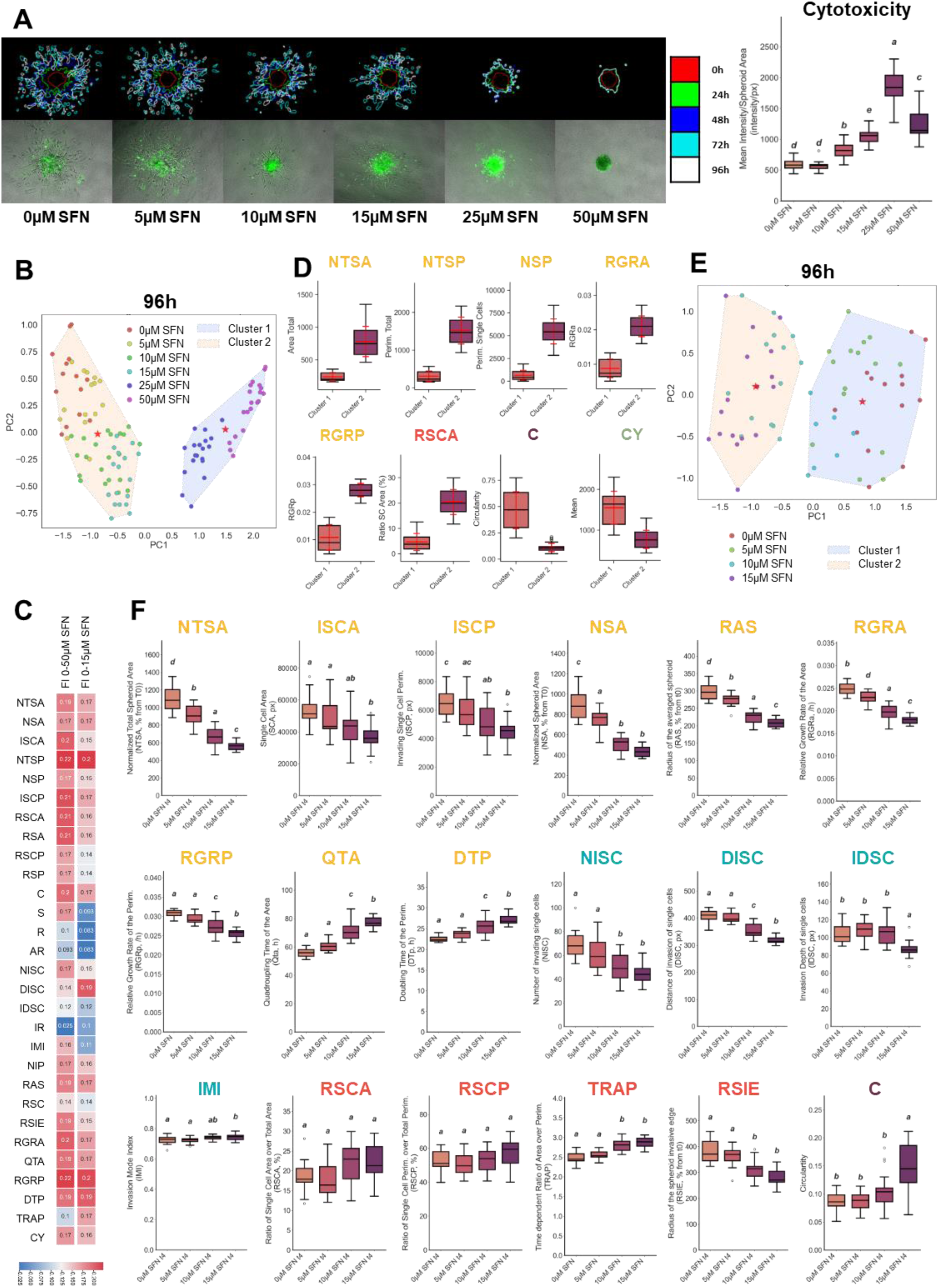
Case study 2 Sorafenib dose-dependent effects in SNU423. (A) SImBA segmentation outlines of representative SNU423 spheroids treated with 0-50µM SFN for 96h (top); SYTOX-Green cytotoxicity staining overlay on raw images (bottom); Cytotoxicity condition-boxplot. (B) k-means clustering after 96h reveals a dominant toxicity driven cluster. (C) FI-ranking before and after removal of the high-dose SFN cluster. (D) k-means Cluster-boxplots after 96h. (E) PCA-rerun with k-means clustering after removing 25 and 50µM. (F) Condition boxplots of selected features at 96h for SFN concentrations (0–15µM). Boxplots show median and IQR; compact letter displays indicate statistically non-significant groups (ANOVA + Tukey or Kruskal–Wallis + Dunn/BYK based on normality).

**Figure 6.**
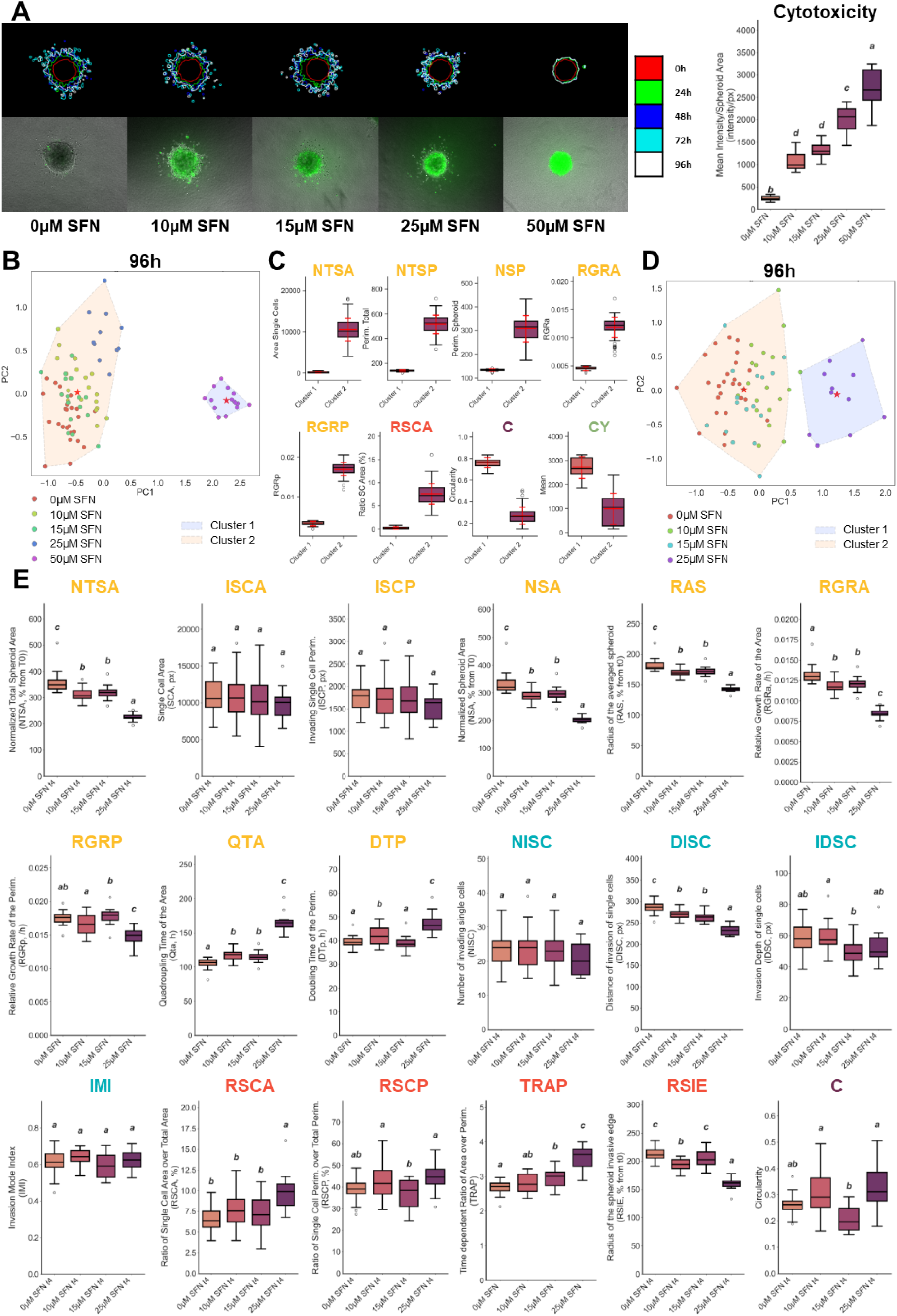
Case study 2 Sorafenib dose-dependent effects in Hep3B. (A) SImBA segmentation outlines of representative Hep3B spheroids treated with 0-50µM SFN for 96h (top); SYTOX-Green cytotoxicity staining overlay on raw images (bottom); Cytotoxicity condition-boxplot. (B) k-means clustering after 96h reveals a dominant toxicity driven cluster. (C) k-means Cluster-boxplots after 96h. (D) PCA-rerun with k-means clustering after removing 50µM. (E) Condition boxplots of selected features at 96h for SFN concentrations (0–25µM). Boxplots show median and IQR; compact letter displays indicate statistically non-significant groups (ANOVA + Tukey or Kruskal–Wallis + Dunn/BYK based on normality).

As expected, SFN induced a dose-dependent cytotoxicity increase after 96h in both SNU423 and Hep3B spheroids (Figure 5A, 6A). PCA-clustering at this timepoint, further showed that high-dose SFN conditions dominated the global phenotypic structure in both datasets. In SNU423 spheroids (Figure 5B), 25-50µM SFN separated from lower doses (5-15µM), whereas in Hep3B spheroids (Figure 6B), the strongest phenotypic separation was observed for 50µM SFN. Here, 25µM SFN clustered with low-dose conditions, indicating that, while cytotoxicity for both cell lines increases dose dependently, SNU423 spheroids are more sensitive to SFN induced toxicity. In the SNU423 spheroids, FI-ranking (Figure 5C) unsurprisingly characterizes this high-dose cluster with a widespread phenotypic shift affecting multiple features, including a.o. total spheroid area and perimeter (NTSA, NTSP), spheroid perimeter (NSP), relative area and perimeter growth rates (RGRA, RGRP), ratio of single cell to total spheroid area (RSCA), circularity (C), and cytotoxicity (CY). Cluster-boxplots for these features show clear reduction in size and growth, invasiveness, lower single-cell area ratio and increased circularity in the high toxicity, high-dose SFN cluster for SNU423 (Figure 5D). For Hep3B spheroids, this same trend of strong decrease of invasion and growth in the high-dose SFN treated spheroids is observable (Figure 6C). Simply put, and consistent with the representative SImBA-generated spheroid outlines and cytotoxicity staining images (Figure 5A, 6A), the data indicate complete inhibition of spheroid growth and invasion coinciding with (or secondary to) maximal toxicity in spheroids treated with 25-50µM SFN (SNU423) or 50µM (Hep3B).

Beyond this apparent clear-cut result, we demonstrate that a more detailed SiQuAl-based analysis of the low-dose SFN range can be highly advantageous. To this end, SiQuAl was rerun while focusing specifically on the lower SFN doses. In this rerun, PCA-clustering of the SNU423 spheroids now separated the remaining conditions into two distinct dose ranges, 0-5µM and 10-15µM SFN, consistent with a gradual, dose-dependent phenotypic response (Figure 5E). For Hep3B spheroids, two clusters were also detected in the rerun, however, this did not reflect a gradual dose-dependent separation across the full dose range. Instead, clustering was driven by the 25µM condition, whereas 0-15µM treated spheroids remained grouped together, indicating limited phenotypic resolution among the lower SFN doses (Figure 6D). Below, we show that SiQuAl allows to couple this to essential differences in underlying biology.

For SNU423, FI evaluation of run 2 PCA-clustering again revealed changes across multiple parameters (Figure 5B), as demonstrated by the condition-boxplots (Figure 5F). Predictably, increasing SFN concentrations were associated with dose-dependent decreases in total and spheroid area (NTSA, NSA), in RGRA, RGRP, in the radius of the averaged spheroid and invasive edge (RAS, RSIE), the distance and depth of single-cell invasion (DISC, IDSC), and in the number, area and perimeter of invading single cells (NISC, ISCA, ISCP). Equally predictable, area quadrupling time, perimeter doubling time and spheroid circularity (QTA, DTP, C) dose-dependently increased (Figure 5F). Slight TRAP increase shows that area growth decreased slightly faster than perimeter growth. Stable ratios of single-cell versus total area and perimeter (RSCA, RSCP) and IMI values indicate that SFN mainly reduced the extent of invasion without markedly changing the invasion mode. Together, this data in the SNU423 spheroids reflects an anticipated, classical toxicity-driven dose-dependent response where increasing cytotoxicity progressively limits spheroid growth, coinciding with an anti-invasive effect. This is expected as growth-inhibited spheroid cells also lose the capacity to invade. The gradual, dose-dependent decrease/increase of multiple features is even more readily apparent in the parallel coordinates plot (Figure S4A).

As noted before, despite the dose-dependency of the cytotoxicity feature (Figure 6A), the rerun 2 for Hep3B spheroids revealed a different lower-dose clustering pattern. After exclusion of 50µM SFN, 25µM treated spheroids formed a single cluster, while spheroids treated with 0-15µM SFN remained in 1 cluster (Figure 6D). Moreover, the Hep3B spheroids did not show a corresponding dose-dependent suppression of growth and invasion (Figure 6F) as seen for the SNU423 (Figure 5F). Although 25µM SFN reduced spheroid area and growth (NTSA, NSA, RGRA, QTA, RGRP, DTP), the radii RAS and RSIE, and DISC, the overall dose-dependency for these features across lower SFN doses is not in line with the gradual, dose-dependent increase of cytotoxicity (Figure 6E versus 6A) and sharply contrasts the SNU423 phenotype (Figure 5F). In addition, multiple (single cell) invasion-associated features remained unchanged across the 0-25µM SFN range or even became relatively more pronounced: NISC did not dose-dependently reduce, RSCA, RSCP increased (Figure 6E). Moreover, also TRAP substantially increased more than in SNU423, indicating perimeter growth was much less impaired than area growth, implying that invasive outgrowth was maintained despite a reduced spheroid bulk. Similarly, although the absolute distance of invading cells (DISC) from the spheroid center decreased, the invasive depth of single cells (IDSC) relative to the spheroid edge remained stable, showing that cells continued to invade comparable distances in relation to smaller spheroid size. Crucially, across the 0-15µM range, little or no significant changes are noticeable (Figure 6E). The parallel-coordinate plots are useful to highlights the divergent response, with substantial overlap between lower-dose conditions in Hep3B in contrast to the SNU423 MCTS (Figure S4). Together, these findings indicate that Hep3B spheroids respond fundamentally differently from SNU423. Higher SFN concentrations increased toxicity and spheroid size is impacted at these high SFN doses for both cell lines, but invasion is not suppressed in the same manner. Combined feature analysis points to sustained invasive activity for HEP3B spheroids irrespective of increasing toxicity. This suggests not only that Hep3B spheroids are less SFN sensitive, but may indicate the presence of a SFN-resistant invasive subpopulation.

This case study shows that, by extracting and integrating a broad panel of quantitative features, SImBA-SiQuAl enables physiologically relevant evaluation of drug responses and reveals phenotypic differences that would be entirely missed in a classical analysis.

## DISCUSSION

Since spheroid and organoid 3D micro-tumour models recapitulate more faithfully in situ tumour heterogeneity, spatial organisation and invasive behaviour than 2D cultures and are substantially more scalable and cost-effective than e.g. rodent models, they have been adopted as effective tools in preclinical and translational cancer research ^10,11,16,19,83,84^. However, the ability to extract and interpret the rich morphological, growth, viability and invasion-related information contained within these complex models remains a major bottleneck. As a result, and despite the high translational potential of these 3D micro-tumour assays, their analysis often addresses a minimal set of rudimentary features, leaving most of their informative potential unexploited^59,62^.

SImBA-SiQuAl directly addresses this major analytical gap by providing an easy-to-use and versatile end-to-end framework designed for quantitative spheroid and organoid image analysis suitable for low- and high-throughput workflows. An essential element to promote SImBA-SiQuAl adoption lies in its accessibility. Both SImBA and SiQuAl can be run as a standalone application, making the workflow easily executable without specialised software or coding expertise. In addition, by covering growth and invasion assays and demonstrating SImBA-based segmentation of both spheroids and organoids, these results support the broad applicability of SImBA-SiQuAl across commonly used 3D microtumour models.

As shown in Table 1, SImBA-SiQuAl addresses important limitations of currently available open-source spheroid-focused analysis tools. Firstly, segmentation quality control is not always implemented within the software or is often performed on an image-per-image basis. This time-consuming manual inspection can reduce optimisation and make analysis of large datasets challenging. SImBA’s Prescreen module, combined with additional artefact removal (via setting the EM), allows evaluating segmentation performance across the entire dataset and all time points at once. This supports dataset-level optimisation of segmentation parameters and helps maximise the use of input images while maintaining robust segmentation across different imaging modalities. To our knowledge, no other toolkit provides an automated global *Prescreen* of this nature. A second distinction is the possibility to integrate viability measurements alongside morphology, growth and invasion metrics. While existing tools primarily focus on morphology and spheroid size, SImBA-SiQuAl incorporates viability as an equal phenotypic dimension. This is of particular interest for high-throughput drug screening. Thirdly, SImBA-SiQuAl supports time-series analysis and calculates additional dynamic features such as relative growth rates and TRAP. Although kinetic readouts are available in some commercial platforms, they are less commonly incorporated into open-source spheroid analysis workflows. Finally, SiQuAl provides an integrated statistical framework that links quantitative feature extraction to comprehensive advanced statistical analysis, guiding users to biological interpretable results. In addition to classical statistical outputs available in some current tools (TASI, SpheroScan), SiQuAl combines multiparametric PCA, clustering and feature-importance analysis into one coherent pipeline. SiQuAl therefore shifts spheroid analysis from single-parameter readouts toward comprehensible global phenotypic profiling.

This phenomics-oriented application is particularly relevant and timely in cancer research, where phenomics are increasingly integrated in multi-omics pipelines. Furthermore, clonal tumour heterogeneity is increasingly recognized as a key driver of metastasis, treatment response and resistance and tumour progression in multiple cancer types^66–68^. Identifying aggressive, drug-tolerant and invasive clones is therefore essential for mechanism-informed drug development and personalized (combination) therapy ^85,86^. In this context, morphology, growth and/or invasion patterns of 3D microtumours provide an indispensable functional readout of genomic, transcriptomic and proteomic states. Several studies further show that omics profiles of 3D microtumours recapitulate in situ clonal dynamics, with patient-derived sub-clonal architectures re-emerging in 3D microtumours^87–90^. This highlights a critical need for unbiased, quantitative phenotyping as a bridge between omics and function. SImBA-SiQuAl may therefore provide benefit for dissecting complex phenotypes in 3D-co-cultures^91^, patient-derived (clonal) spheroids or tumour organoids. Both case studies position SImBA-SiQuAl precisely in this gap, by demonstrating multiparametric, data-driven quantification and subtyping of invasive or drug-induced behaviours with greater resolving power than single-metric approaches.

Case 2 clearly demonstrates why SImBA-SiQuAl provides added value. Since 3D microtumours are complex, multifaceted systems, reducing treatment response to rudimentary area- or viability-based endpoints is inherently misleading thereby possibly obscuring or not measuring clinically relevant effects. The detailed SImBA-SiQuAl output identified the Hep3B spheroids are capable of maintaining invasive behaviour despite increased drug-induced toxicity. This supports accumulating evidence that increased invasion and epithelial–mesenchymal transition tightly links to sorafenib treatment resistance in hepatocellular carcinoma^92–95^. Our findings support a model where therapeutic pressure can enrich for cells with invasive drug-resistant phenotypes in specific patients or subclones rather than uniformly suppress overall tumour aggressiveness (here exemplified by SNU423 responses). Crucially, this sustained invasion phenotype would have remained undetected if analysis had been restricted to conventional screening readouts such as spheroid area, diameter or global toxicity alone. For the above reasons we promote SImBA-SiQuAl as a high-throughput, multiparametric 3D microtumour phenomics strategy, that can provide a functional basis for or readout of genomic, transcriptomic and proteomic data and which enables test strategies for pathway-level mechanistic insight into functional tumour behaviour. Moreover, SImBA-SiQuAl provides a practical framework to quantify patient heterogeneity and personalized treatment response^96^ in a physiologically relevant test system, such as (patient-derived) spheroids or tumour organoids.

In conclusion, SImBA-SiQuAl opens possibilities to harness the full informational complexity of tumour spheroids and organoids thereby addressing central challenges in modern 3D cancer research. By coupling segmentation QC, robust and unbiased feature extraction, integrated viability measurements and an extensive statistical analysis suite, the platform provides a high-resolution, physiologically relevant representation of spheroid behaviour. Its accessibility and reproducibility positions it as a scalable tool for drug-induced cancer phenomics and for integration in multi-omics applications aimed at decoding functional tumour heterogeneous behaviour and responses.

### Limitations of the study

We demonstrated applicability to organoid segmentation but we did not yet extend validation to complex organoid assay workflows. Broader implementation in organoid-specific models, including heterogeneous and structurally complex systems, will therefore require further optimization and benchmarking. Furthermore, the current workflow primarily relies on 2D image information and is therefore not yet optimized for full z-stack acquisition, volumetric reconstruction, or true 3D projection-based analysis. As a result, biologically relevant information present along the z-axis may be underrepresented. In addition, fluorescence support is currently limited to endpoint staining and would benefit from broader integration of continuous live/dead staining and other longitudinal fluorescent readouts. This could further strengthen assessment of viability, cytotoxicity, and dynamic treatment responses over time. Finally, the parallel coordinate plots shown in supplement will be implemented in a future update of SiQuAl. Addressing these points will be important to further expand the biological scope and translational relevance of the SImBA-SiQuAl framework. Given the open-source nature, addressing these could be community-driven or - supported.

## Supporting information

Supplementary Materials: Additional SImBA-SiQuAl workflow information(1, 1.1, 1.2, 2) , Figures S1-S4; Table S1-S5

## RESOURCE AVAILABILITY

### Lead contact

Requests for further information should be directed to the lead contact, Elias Van De Vijver

## Materials availability

No new unique reagents were generated.

## Data and code availability

- Case study spheroid images and a full SiQuAl analysis of each case are available at Figshare: [10.6084/m9.figshare.31268722] and will be accessible as of the date of publication.
- Example datasets and an accompanying SImBA-SiQuAl manual are available at Figshare at [10.6084/m9.figshare.30639479] and will be accessible as of the date of publication.
- All original code for SImBA-SiQuAl is available at Figshare at [10.6084/m9.figshare.31268728] and will be accessible as of the date of publication.
- Additional information required to reanalyse the data reported in this study will be made available upon reasonable request.

## ACKNOWLEDGMENTS

EVDV was supported by the Research Foundation – Flanders (FWO1SE0722N) and UGent Bijzonder Onderzoeksfonds (BOF.BAF.2024.0285.01, from CA). HVV was supported by FWO senior research grant (FWO1801721N).

## AUTHOR CONTRIBUTIONS

Conceptualization, EVDV, MVT.; methodology, EVDV, KD, AVA; Investigation, EVDV, KD, AVA, MVT.; writing-original draft, EVDV; writing-review & editing, MVT, KD, AVA, HVV, CA, EVDV; funding acquisition, MVT, HVV, CA; supervision, MVT

## SUPPLEMENTAL INFORMATION

**Supplementary Materials: Additional SImBA-SiQuAl workflow information(1, 1.1, 1.2, 2) , Figures S1–S4; Table S1-S5**

## STAR★METHODS

### KEY RESOURCES TABLE

| REAGENT OR RESOURCE | SOURCE | IDENTIFIER |
| --- | --- | --- |
| Antibodies |  |  |
| / |  |  |
| Bacterial and virus strains |  |  |
| / |  |  |
| Biological samples |  |  |
| / |  |  |
| Chemicals, peptides, and recombinant proteins |  |  |
| Hepatocyte Growth Factor human | Sigma-Aldrich | Cat# h5791 |
| Sorafenib tosylate | MedChemExpress | Cat# HY-10201A |
| Collagen I, rat tail, 3,5mg/ml | Corning | Cat# 354236 |
| SYTOX™ Green Dead Cell Stain | Molecular Probes™ | Cat# 11310972 |
| Critical commercial assays |  |  |
| / |  |  |
| Deposited data |  |  |
| Case study data + Results | This paper | https://doi.org/10.6084/m9.figshare.31268722 |
| Example datasets + manual | This paper | https://doi.org/10.6084/m9.figshare.30639479 |
| Experimental models: Cell lines |  |  |
| Hep3B2.1-7 (Hep3B) | Cytion | CVCL_0326 |
| Huh-7 | ATCC | CVCL_0336 |
| SNU-423 | ATCC | CVCL_0366 |
| Hela | ATCC | CVCL_0030 |
| Hepa 1-6 | ATCC | CVCL_0327 |
| MDA-MB-231 | ATCC | CVCL_0062 |
| HT-1080 | ATCC | CVCL_0317 |
| SK-MEL-28 | ATCC | CVCL_0526 |
| Experimental models: Organisms/strains |  |  |
| / |  |  |
| Oligonucleotides |  |  |
| / |  |  |
| Recombinant DNA |  |  |
| / |  |  |
| Software and algorithms |  |  |
| SimBA-SiQuAl | This paper | https://doi.org/10.6084/m9.figshare.31268728 |
| CellSens | Olympus/Evident Scientific | https://evidentscientific.com/en/products/software/cellsens |
| Python | Python Software Foundation (PSF) | https://www.python.org/downloads/ |
| FIJI | Shindelin et al, 2012 <sup>97</sup> | https://imagej.net/software/fiji/downloads |
| Other |  |  |
| / |  |  |

### EXPERIMENTAL MODEL AND STUDY PARTICIPANT DETAILS

#### METHOD DETAILS

##### Cell lines

Spheroids of 8 cell lines originating from variable tumour origin were examined. Huh-7 (CVCL_0336) and SNU-423 (CVCL_0366) cells were cultured in RPMI-1640 medium (1X) + GlutaMax TM-I (Gibco) supplemented with 1% (v/v) sodium pyruvate (Gibco), 10% (v/v) foetal calf serum (FCS, TICO Europe) and 1% (v/v) Antibiotic-Antimycotic (AA; Gibco). Hep3B2.1-7 (CVCL_0326), Hela (CVCL_0030), Hepa 1-6 (CVCL_0327), MDA-MB-231 (CVCL_0062), HT-1080 (CVCL_0317) and SK-MEL-28 (CVCL_0526) were cultured in Dulbecco’s Modified Eagle Medium (DMEM) (1X) + GlutaMax TM-I (Gibco) supplemented with 10% (v/v) FCS (TICO Europe) and 1% (v/v) AA (Gibco). Cell cultures were passaged 1 to 3 times a week depending on the cell line using standard procedures. Cells were maintained in a humidified incubator at 37°C at 5% CO_2_.

##### Spheroid generation

All spheroids were generated using the hanging drop method to facilitate ECM embedding in later stages. Accordingly, cells were trypsinized and resuspended in the appropriate complete culture medium into a single cell suspension (CS; 100.000 cells/ml). To increase the viscosity of the medium, methylcellulose (MC; Sigma Aldrich) (12 mg/ml in medium containing 2% v/v AA) was added in a 4:1 CS-MC mix to obtain a final cell concentration of 80.000 cells/ml. Of this mix, drops of 25µl containing 2000 cells were pipetted onto the hydrophobic lid of a square plastic cell culture disc (Nunc, 240835) using a multichannel pipette. 10 ml of phosphate buffered saline (PBS) (Gibco) was added to the bottom dish to avoid droplet evaporation. The droplets were placed in a humidified incubator (37°C, 5% CO2) until solid spheroids were detectable under the microscope (LEICA MZ7.5): 24h for Hep3B2.1-7, Huh-7, SNU-423, Hela, Hepa 1-6 and 72h for MDA-MB-231, HT-1080, SK-MEL-28.

##### Collagen embedded spheroid invasion assay

Spheroids (± 15/well) were embedded in a 3D collagen type I extracellular matrix (CECM) in a 12-well plate (Corning) according to Van Troys et al^98^. The collagen solution consisted of precooled Hank’s Balanced Salt Solution (HBSS, GIBCO), 0.72xMEM (GIBCO, from 10XMEM), 18 mM NaHCO_3_ (from 0.25M), 1/5 volume complete growth medium (+10% FBS, + 1% AA), 1mg/mL collagen (CORNING COLLAGEN I, RAT TAIL, 3,5mg/ml) and 13 mM NaOH (from 1M stock) added in this respective order and prepared on ice to prevent polymerization. The 12-well plate was kept at 37°C for least an hour at before use. The temperature of the well plate was kept constant during the entire process of embedding using a heating plate set at 37°C. The 3D CECM consisted of 2 layers. In the preheated 12 well plate, a bottom layer (350 µl) was pipetted per well and distributed evenly on the well bottom by gently tapping the sides of the well plate. The bottom layer was left to polymerize for 20 minutes and has as purpose to prevent subsequently added spheroids from settling and adhering to the bottom of the well plate. During this polymerization of the bottom layer, 15 spheroids were manually picked (using a p1000 pipet tip) under a binocular microscope (LEICA MZ7.5) and transferred to a 2ml Eppendorf tube. Subsequently, spheroids were gently washed with growth medium to remove MC rich medium. Spheroids were allowed to settle in the bottom of the tube; medium was removed and spheroids were resuspended gently in 360µl cold CECM and pipetted dropwise onto the bottom layer ensuring they are well separated. The top layer is left to polymerise on the heat plate for an additional 20 minutes to prevent the spheroids from moving around while transferring the well plate to the incubator. Subsequently, the well plate was left in the incubator for an additional 40 minutes. After this polymerisation step, 750 µl of culture medium was added to each well. T_0_ images were captured at this stage. After this, 50 ng/ml hepatocyte growth factor (HGF, Sigma) was added to each well to induce invasion into the CECM. Where mentioned, the culture medium contained Sorafenib tosylate (Bay 43-9006 tosylate; MedchemExpress) at te indicated dose. Images were acquired using phase-contrast imaging (Olympus IX 81, Olympus CellSens imaging software) at different time points (0-48 or 0-96h, as indicated) using a 10x objective. In short, the well plate was transferred to an Okolab microscope incubator at 37°C. Using the CellSens imaging software, spheroid positions were manually selected, saved in a position list and imaged with autofocus (range of 200nm). Each spheroid was given a name based on the plate letter, well ID, spheroid number and timepoint (e.g. AA01_01 t1: first plate, well A1, first spheroid, second time point). Spheroids were imaged every 24h. At the final time point, spheroids were stained using SYTOX-Green (Molecular Probes) (200nM; 30 minutes incubation) to assess toxicity. At this time point, spheroids were imaged twice: phase-contrast followed by fluorescent imaging. Fluorescent images of the latter were named according to spheroid ID (AA01_01) and saved in a “FLUO” subfolder.

##### Use of Public Dataset

Publicly available spheroid images were obtained from the Spheroid Light Microscopy Image Atlas (SLiMIA)^82^. Images from five different microscope systems (Axiovert200M, Cytation5, IncucyteZOOM, OlympusIX05, Opera Phenix) were selected based on overall image quality containing an in-focus spheroid. All raw images were cropped to remove embedded scalebars and subjected to the best performing segmentation method in SImBA as outlined in Table S4. Segmentation methods was kept consistent for each cell line.

### QUANTIFICATION AND STATISTICAL ANALYSIS

Statistical analysis was performed according to the described protocol discussed in the results and Supplementary Materials. In short, for all features, normality was checked using a Shapiro Wilk test. Based on this normality, at all timepoints except t_0_, SiQuAl automatically performs either one-way ANOVA with Tukey post-hoc testing or Kruskal-Wallis with Dunn post-hoc with BYK p-value correction. Boxplots represent the median with interquartile range (IQR). Statistical significance is shown as compact letter display, where non-significant groups bear the same letter. The pairwise statistical comparisons between experimental conditions are also visualized using non-significance networks as well as p-value heatmaps (not shown in this manuscript, available at: https://doi.org/10.6084/m9.figshare.31268722), *p<0.05, **p< 0.01, ***p<0.001, ****p<0.0001. The total number of spheroids included in both case studies is described in detail in Table S5.

### ADDITIONAL RESOURCES

As mentioned, all raw and analysed case study material is provided at https://doi.org/10.6084/m9.figshare.31268722.

Additionally, example datasets including full SImBA-SiQuAl analysis and a SImBA-SiQuAl manual are available at: https://doi.org/10.6084/m9.figshare.30639479.

Code of both SImBA and SiQuAl are available at: https://doi.org/10.6084/m9.figshare.31268728.

