## Supplementary Materials: Additional SImBA-SiQuAl workflow information(1, 1.1, 1.2, 2) , Figures S1-S4; Table S1-S5 for "SImBA-SiQuAl: advancing high-content high-throughput phenotypic profiling of 3D microtumours"

### ADDITIONAL SIMBA-SIQUAL WORKFLOW INFORMATION

#### 1. *SImBA-SiQuAl: complete workflow from raw spheroid image series to in-depth biological insight*

In this supplementary material, we provide additional information in the form of more detailed descriptions on specific steps depicted in main text Figure 1 either on the SImBA part (1.1. below) or on the SiQuAl part (1.2. below). The numbered steps in main text Figure 1 are consequently also used as reference here (e.g. Figure 1, step X).

In 2. below, we additionally provide a 'results walkthrough' explaining how to explore and fully exploit in the best/optimal manner the extensive output of SImBA-SiQuAl. This supplements the briefer description in the main text and associated main text Figure 2B.

To become familiar with SImBA-SiQuAl two example data sets are made available on Figshare: https://doi.org/10.6084/m9.figshare.30639479. The example data sets are smaller versions (i.e. with a more limited number of conditions) of the data used for Case study 1 and 2 in the main text as detailed in the "README" file available with the example datasets. The example data can be used by first users to run the software using the guides through the SImBA-SiQuAl analysis (see README). Additionally, with the example datasets, the output of a complete SImBA-SiQuAl run is available (see README).

Before the user can initiate the workflow (Figure 1 step 1), the image directory with the raw image data must conform requirements as described in Figure S1A. This concerns names of (sub)folders and corresponding input image files (.TIFF) and the specific directory layout of (sub)folders in the main directory on which the software (i.e. the SImBA-part) runs.

##### 1.1. *SImBA: Spheroid Image Batch Analysis*

- SImBA is a FIJI macro and can be run as such in FIJI either via *Plugins* ‣ *Macros* ‣ *Run*… or by opening the script through *File* ‣ *New* ‣ *Script*… and executing Run (Figure 1, step 2). We however recommend to run SImBA via the executable created specifically to run SImBA. This executable will run the _SImBA.ijm macro in a suitable version of FIJI/Imagej to avoid any dependency issues. This is further explained in the README file provided with the example datasets.

- Spheroid growth or Spheroid invasion? As specified in the main text, the first selection the user makes is to choose between a "*Growth run*" (Figure 1, step 3A) or an "*Invasion run*" (step 3B). SImBA indeed offers analysis of spheroid images from two commonly used distinct assay formats:

  (i) A spheroid growth assay where spheroids are imaged while cultured in growth medium, usually in the ultralow attachment (ULA) multiwell plates in which they have been generated. Here the phenotype of interest is the spheroid shape, morphology and expansion by growth.

  (ii) A spheroid invasion assay where spheroids are imaged while embedded in a 3D-hydrogel (e.g. collagen or Matrigel). Here, (invasive) growth of the spheroid can co-occur with other phenotypes such as single cell invasion or invasive strand formation.

  In the spheroid invasion format, it is of the utmost importance to quantify features of both the spheroid and the invading areas (single or grouped cells) that in time populate the extracellular matrix and are distant from the spheroid mass. On the contrary, for a spheroid growth assay, any cells outside of the growing spheroid should not be considered in the quantification. Images of spheroids in medium are usually generated from a single cell suspension and indeed often contain 'loose' cells or debris in the medium (Figure S1B). Therefore, in a growth assay, only the spheroid is the object of interest and the (remaining) single cells still present in the image usually have no biological relevance. The two pipelines In SImBA, despite being very similar, serve these two different biological assay setups in the following manner: during image processing in a *Growth run* (Figure 1, step 3A, Figure S1B), only the largest segmented area (corresponding to the spheroid) is retained in the mask and considered during subsequent quantification and feature extraction of the segmentation masks, as indicated in green in Figure S1B (top two rows). In contrast, in an *Invasion Run* **(**(Figure 1, step 3A), all segmented areas are expected to contribute to the final SImBA output and thus used throughout the analysis (Figure S1B, bottom two rows). We note that

this choice strongly expands SImBA-SiQuAl applicability. As additional benefit, the settings for 'growth only', also enables spheroid analysis for datasets with numerous debris or artefacts in the medium/matrix that otherwise would be entirely unusable for quantitative analysis.
For Figure 1, step 4 where the user defines the "*main directory"* to analyse we recommend to perform the analysis on a copy of the raw data.

- Running a *Prescreen* (Figure 1, step 5):
  Running a *Prescreen* is optional in SImBA, but has three main advantages:

  A) Testing for the optimal segmentation method (Figure 1 step 5A *(i)*): Up to five different segmentation methods are available in SImBA and can be tested using the *Prescreen* module to identify which best delineates spheroids in your image set. Iterative *Prescreen* runs with different settings will not affect downstream analysis as they are run in temporary directories and thus do not affect the original image data.
  The five segmentation options are (i) ***Default*** segmentation (*'No Auto Threshold'; NAT*) which is performant in the majority of cases by combining multiple smoothing steps with the *Find edges* plugin and *Gaussian blur* to delineate the spheroid from the background; (ii) ***Find edges (FE)*** relies heavily on the built in *Find edges* plugin to, after enhancing the contrast of the image prior to a smoothing step, use as input of auto-thresholding to allow to delineate the spheroid from the background. This method works very well on strongly invading spheroids; however, segmentation accuracy decreases when the background (ECM) is very light compared to the high contrast (dark) spheroid; (iii) ***Auto threshold (AT).*** This method starts by enhancing the contrast of the image prior to a sharpening step. Subsequently background subtraction is performed followed by a Gaussian blur using a user defined Sigma value (see below). This is followed by auto-thresholding to delineate the spheroid from the background. Although this method is successful on almost all spheroids irrespective of contrast or debris in the image, this is a very stringent segmentation method that should only be used as "last resort" as it often overlooks invading single cells and reduces protruding edges; (iv) ***Double Blur (DB)***. This method combines two Gaussian blur steps interspersed with a Find Edges step without any pre-processing of the image (i.e. enhancing the contrast or sharpening/smoothing). This segmentation method performs comparably to the default (NAT) segmentation approach. Because it applies two consecutive blurring steps, it is less prone to over-segmenting small background areas. However, this additional smoothing can also slightly reduce (the detection of) invading areas; (v) ***Fluorescence (FL)*** segmentation. While most segmentation methods are focussed on segmenting brightfield and phase contrast images, the segmentation of fluorescent images is more obvious due to the lack of contrast fluctuations and or ECM structure variations. By enhancing the contrast and smoothening the image, auto-threshold is sufficient to segment both the spheroid and its invading cells.

  B) Sigma tweaking (Figure 1, step 5A*(ii)*): The *sigma* parameter controls the variance of the Gaussian kernel (i.e. the extent of Gaussian blur). A higher sigma causes stronger smoothing and vice versa. The ability to easily test this value in advance across all test conditions is a clear advantage in the *Prescreen* module, as it allows users to balance the risk of background over-segmentation at low sigma values, where small irregularities may be falsely identified as invading areas, against excessive smoothing at high sigma values, where smaller invading regions are no longer distinguished from the background and perimeter detail is lost.

  C) Defining the Error margin (EM) (Figure 1, step 5B*)*). The EM is a multiplier of the averaged spheroid radius (RAS, radius of the smallest circle containing all area of the total spheroid). Areas outside this *radius* × *EM* are removed. This setting of EM thus allows to filter 'debris' (a term here used to cover non-spheroid, non-invasive objects including cell (fragments), dust, matrix artefacts, etc.) in the spheroid periphery that is unrelated to the spheroid of interest. This prevents overestimation of invasion as artefacts in the image periphery would otherwise be considered as "invasive cells". The EM is particularly important when multiple spheroids are cultured in a single well, and when e.g. a second spheroid appears in the same field of view (as illustrated by the green arrows in Figure S1C). The choice of EM is facilitated by the coloured concentric circles (with increasing EM-value) displayed on the *Prescreen* montages (Figure S1C).

The selection of an appropriate segmentation method, Sigma- and EM value are facilitated greatly as the *Prescreen* segmentation results are presented as a handy large montage containing all spheroids per timepoint. Figure S1C contains only a subset of the *Prescreen* result; full montage examples can be assessed in Table S2, 9-13. SImBA prompts the user to visually inspect the *Prescreen* output by scrolling through the montage stack of *Prescreen* montages in time (Figure 1, step 5B*)*. At this stage, it is possible to remove problematic spheroids from the main directory if needed[1].

The *Prescreen* step is highly recommended, yet optional. If the *Prescreen* is skipped (Figure 1 step 5, red dotted arrow), SImBA will use the _*default*_ segmentation method. The EM should then be set at 3000. This "arbitrarily large" EM can also be used if no debris needs to be filtered from the edges of the image. The EM in a GROWTH run is of less relevance as no additional areas are retained after segmentation.
The *Prescreen* process can be used in an iterative manner: the user can either decide to re-run the *Prescreen* to assess different parameter settings Figure 1, step 5D*)* or continue SImBA with the last tested segmentation settings if satisfactory.

Taken together, the advantages of the SImBA *Prescreen* address a key need familiar to all users working with spheroid imaging: the ability to exclude problematic images in a standardized and reproducible manner. Importantly, this exclusion is based on the actual segmentation outcome rather than on subjective preselection prior to analysis, thereby improving consistency across experiments and users.

- _Fully automated SIMBA run_: Before the automated run, the user is prompted to define the experimental and analysis settings including number of timepoints, imaging interval (constant or variable) and chosen EM and sigma values. The user also indicates whether visual outputs should be generated, and whether quantification of cytotoxicity from fluorescent images is applicable (Figure 1, step 6*)*. For very large datasets, comprising numerous spheroids and/or time points, we recommend generating visual outputs only for a representative subset in order to avoid excessive storage requirements.
  As detailed in main text, SImBA segments all included spheroids fully automatically (Figure 1, step 7-11*)* and extracts for all spheroids at each time point the numerical values for the full list of parameters (main text Figure 2, Table S1). We note here additionally that these values are saved in the *Measurements* folder (in 8 CSV files, see Table S2,1-8, see also guided Examples). The *Measurements* folder is input for the SiQuAl run. SImBA also generates and saves visual output (if selected). Six types of visual SImBA output (for details see Figure S1D and legend) are stored in the individual spheroid folders as detailed Table S2, 45-70, example images included). The final *Prescreen* montages can also be saved (Figure 1, step 12; Table S2, 9-13*).* All individual steps during analysis are reported in the Log window for reproducibility and review in case of error. This log window is saved in a timestamped (yyyy-mm-dd_hh-mm-ss) text file stored in the Measurements folder.

#### *1.2. SiQuAl: SImBA Quantitative Output Analysis*

As descripted in the main text, SiQuAl integrates seamlessly downstream of SImBA. It can be launched either by running the main script in a terminal, provided Python 3 is installed on the user's desktop, from a Python editor such as VS Code, or via the standalone executable (windows) (detailed in README). The module consists of three scripts: the *SiQuAl.py* driver script, which imports the scripts *Classes.py* and *StatisticalAnalysis.py*. In practice, if not running via the executable, running the *SiQuAl.py* driver script is

[1] NOTE: a future update will provide a selection list to easily include or exclude spheroids directly from a checkable list.

sufficient if the other scripts are stored in the same directory. For repeated runs of the same experiment or very large, multi-condition datasets it can be more convenient to run SiQuAl from a dedicated configuration script in which all conditions are hard-coded as described for the *Example Datasets 1 and 2* in the README file (available at: https://doi.org/10.6084/m9.figshare.30639479).

Upon launch, SiQuAl will first remind the user to save results from a previous SiQuAl run, to avoid accidental overwriting. Ensure such outputs are moved to a directory outside the "*Measurements*" folder before proceeding. Similar as to SImBA, before automated run, the user is asked to define the experiment (Figure 1, step 15) This includes the path to the SImBA Measurements folder as input, whether cytotoxicity was measured, the experiment size (number of spheroids per condition and timepoints) and condition names and whether quantile–quantile (QQ) normality plots should be plotted (time consuming). In contrast to SImBA, there is no need to work on a copy of the numerical data as the original measurement (.csv) files are never altered. After this, the user is again asked whether SiQuAl may clear output folders from a prior analysis as an additional check. This should only be confirmed if this is the first run or if previous outputs have been backed up elsewhere.
Once the information above is provided, SiQuAl will first structure and normalized the input .csv data to the reference (t0) timepoint for each spheroid where applicable (Figure 1, step 15) (see Table S1 for normalized feature definitions). The csv-files are transformed by SiQuAl into single csv/xlsx files containing all spheroid values of 1 parameter for 1 timepoint. The rows in these newly generated csv-files represent individual spheroids, whereas the columns are sorted conditions, normalized to timepoint t0 if applicable. All these intermediate structured and/or normalized files can be found in the "*Output*" folder (Table S3).

As detailed in main text and Figure 1, SiQuAl then performs two complementary layers of analysis. Runtimes for both the statistical- and PCA workflows are individually reported, as well as the runtime of the complete SiQuAl run. We here provide additional info for each analysis layer not detailed in the main text.

- Comprehensive statistical evaluation per-condition/per feature/per time-point (Figure 1 step 16A-D).
    - As overview output tool intended to provide a general overview of the dataset, plots are provided per feature in which all timepoints and conditions are grouped within a single plot (step 16A), displayed either as a bar plot or as a bar plot with scatter overlay to visualize data variation (description: Table S3, row 45-47). In the current version this is mainly considered as a working tool for users.
    - Normality assessment of all feature data at each timepoint (step 16B). The results are reported in the Normality.txt file in the Stats/Normality folder (Table S3) and can optionally also be visualized through QQ plots for each feature across all timepoints, although this additional plotting step can be time-consuming and storage-intensive. Based on the normality assessment, SiQuAl subsequently performs the appropriate parametric or non-parametric statistical tests (ANOVA or Kruskal–Wallis, respectively), followed by the corresponding post hoc multiple-testing procedure (Tukey or Dunn test with BYK correction, respectively) (step 16C). Results are stored in de output *Stats* and *Plots* folders as detailed in Table S3.
    - Output: These results are reported through condition-boxplots per timepoint with compact letter display, p-value matrices and heatmaps, and non-significance networks (step 16D, available at https://doi.org/10.6084/m9.figshare.31268722) and *Plots* folders as detailed in Table S3.
- Multivariate analysis by means of principal component analysis (PCA) and clustering (Figure 1 steps 16A-D)
    - Clustering method. The user selects the clustering method, either k-means or DBSCAN, and defines a feature-importance threshold for secondary PCA refinement. K-means clustering will divide observations into a predefined number of groups (k) by minimizing the distance between each observation and the centre of its assigned cluster. The Calinski–Harabasz index is used to determine the optimal number of groups automatically. DBSCAN, in contrast, identifies clusters based on local point density without requiring a predefined number of clusters, allowing it to detect irregularly shaped groups and classify sparsely positioned observations as noise or outliers. We recommend k-means as the default option, whereas DBSCAN may be preferable for PCA distributions containing outliers or several compact, dense groups. However, both can easily be performed consecutively, as they may yield complementary insights. Because DBSCAN is sensitive to differences in cluster density, it can occur that it combines groups with

varying density that would appear as obvious clusters to the user, in which case k-means is again preferred.

- PCA/cluster-analyses and graphical output, feature importance calculation and ranking: Two consecutive PCAs are performed using the selected clustering method. The first PCA includes all features and should be used as main SiQuAl result, whereas the second, refined PCA excludes features below a user-defined importance threshold, determined from the initial PCA. Based on experience, a default threshold of 0.15 removes low-impact features while retaining most informative parameters. Refined PCA and clustering outputs are saved with the prefix NEW_ (see Table S3). If the initial PCA already shows clear separation, inspection of the refined PCA is optional; if not, it may reveal more subtle group differences (see case study 1). Importantly, the absence of differential clustering in the initial PCA is itself informative, indicating limited phenotypic divergence between groups. This procedure can be repeated to test alternative feature-importance thresholds or feature selection (see case studies).
- Cluster-boxplots for all parameters and timepoints for both the initial and refined (NEW_) PCA results (Table S3, Figure 2B).

### 2. SImBA-SiQuAl Results Walkthrough

While the output of SImBA is relatively straightforward (different visual outputs and numerical output for features in .csv files), the SiQuAl output is rich and putatively complex to handle for new users. As indicated in the main text, a multitude of parameters is extracted for each spheroid and for every timepoint. This makes inspection of every feature/timepoint individually neither practical nor efficient. Consequently, an all-inclusive, multiparametric data output via PCA and clustering strategies, coupled with insight into key parameters that drive the cluster separation (via feature importance ranking and cluster-boxplots) and linked statistical comparisons of how a specific parameter changes (or not) across conditions via condition-boxplots is needed.
For transparency, when SiQuAl is run via the executable, all print statements normally available in the console are stored in a SiQuAl_log.txt file containing timestamped steps executed during the analysis. Similarly, in case an error occurs, the error statement is also saved in an SiQuAl_error_log.txt file. Below we provide a 'model' guideline on how to systematically explore and exploit the SiQuAl output and extract biologically meaningful conclusions. This supplements main text Figure 2B. The recommended approach starts from a global view.

- To start evaluating a SiQuAl analysis result, the user first assesses whether distinct clusters emerge after the non-prioritized principal component analysis (PCA 1, all features taken into account) using the cluster-PCA plots (either k-means or DBSCAN). It is important to not only focus on the final timepoint, as spheroid (invasion) phenotypes evolve over time and interesting kinetic differences may also be revealed by an evolution of the clustering in time (see case study 1 in main text). Therefore, each timepoint should be assessed independently, while taking into account that clusters may evolve between timepoints.
- When clusters between test conditions are observed, the user subsequently reviews the feature-importance outputs provided as CSV files, heatmaps (using blue-red gradient colouring) and/or feature importance ranked console prints to identify which features most strongly drive cluster separation. The feature-importance table calculates the weighted average of all absolute principal component loadings weighted by their explained variance. In addition, and for a more detailed insight, the PCA-Loadings.csv file reports the correlation between each original parameter and each new principal component, indicating how much each variable contributes to a component. This also indicates whether discrimination of clusters for a specific feature occurs primarily along PC1 (x-axis) or PC2 (y-axis). In the console window, as mentioned above, SiQuAl will also print a ranked list of features with an importance higher than the threshold. This allows users to readily prioritize features at each timepoint over less informative parameters.
- To understand how these prioritized features, contribute to clustering, or distinguish conditions, cluster-boxplots (x-axis = cluster ID) can be interpreted alongside condition-boxplots (x-axis = condition), which also display the results of statistical testing. For a visual representation of the statistical correlations, SiQuAl generates non-significance network plots for each feature at every timepoint. Together, this enables rapid, biologically meaningful identification of the features that

most strongly contribute to the observed clustering and thus to the biological differences between spheroid phenotypes from different test conditions.

- As described above, SiQuAl first calculates feature importance from the PCA including all features and subsequently uses this ranking to perform a second PCA on a refined feature set. Evaluation of this second ("NEW_") PCA is optional when the initial PCA already reveals clear and biologically meaningful clustering. However, when the initial PCA does not show distinct clusters, a likely explanation is the inclusion of too many weakly varying or non-discriminative parameters. In such cases, the NEW_PCA allows the user to reassess potential differences between conditions after reducing noise by excluding low-impact features. If clustering emerges after this refinement step, feature importance should again be interpreted alongside cluster- and condition-based boxplots to identify the parameters driving separation. It is equally important to inspect which features were excluded in the NEW_PCA, as these represent stable, non-discriminatory parameters that may themselves be biologically informative by highlighting spheroid characteristics unaffected by the perturbation employed e.g. drug treatment.
- If the refined PCA still fails to reveal biologically meaningful clusters, possible explanations include a feature-importance threshold that remains too low, inclusion of too many experimental conditions in a single analysis, or a genuine lack of phenotypic differences between conditions. In that case, the user may either increase the threshold or repeat the SiQuAl analysis after excluding selected conditions at the SImBA .csv level to improve contrast. If clustering remains absent after these refinements, the most likely conclusion is that the analysed conditions do not differ sufficiently at the examined timepoint.

By moving from a global, multi-feature view to a focussed, feature-refined view, the user can in this manner convert the broad (SImBA-)SiQuAl output into an interpretable, phenotype-based description to gain clear, biological insight into the dataset.

### SUPPLEMENTARY FIGURES

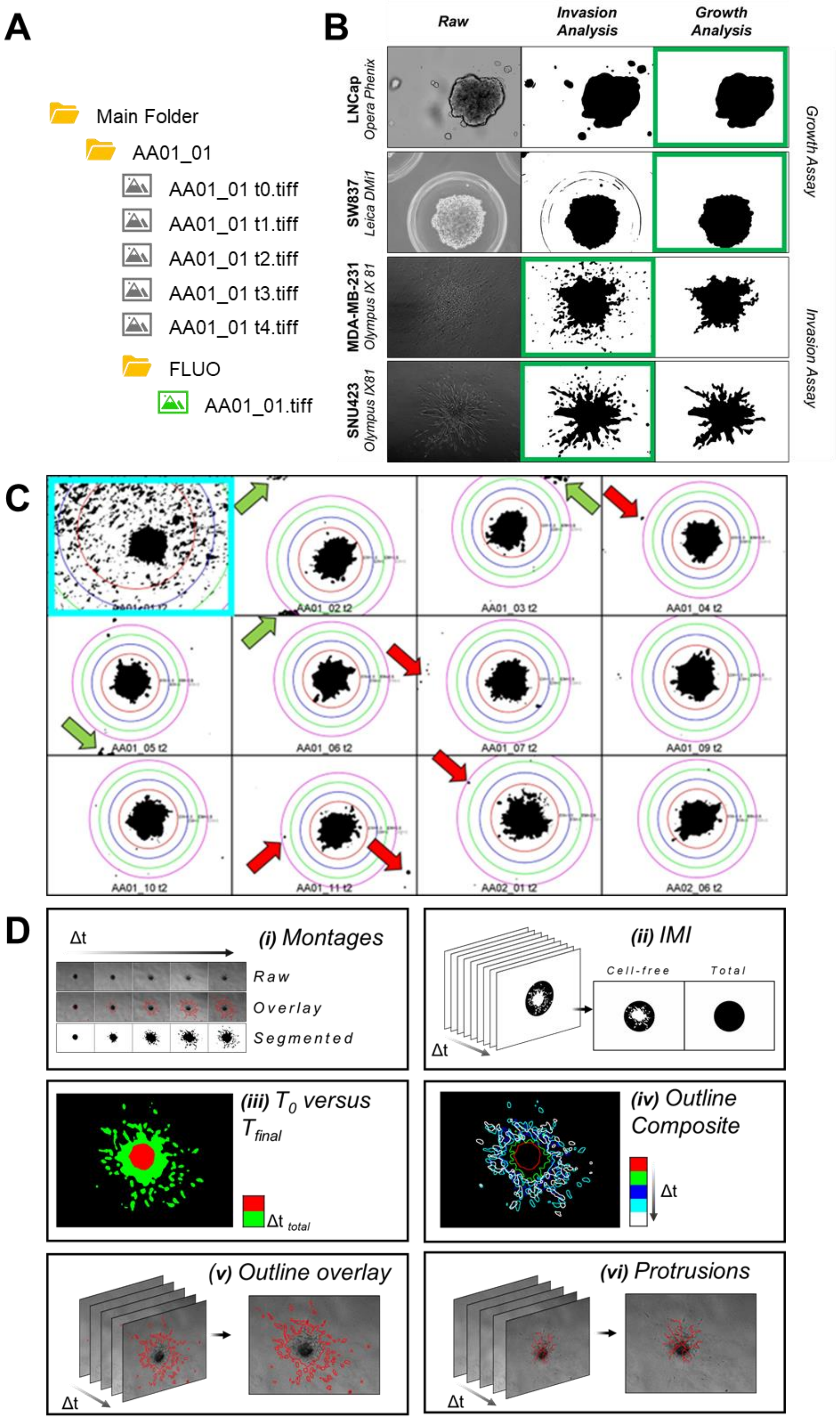

***Figure S1. Details on SImBA directory structure, growth or invasion assay applicability, Prescreen evaluation and the different generated visual outputs.***

(**A**) At the start of the SImBA segmentation run, individual main directory subfolders should contain images of 1 spheroid over multiple timepoints (TP). TP numbering should start with “t0”. Fluorescence viability/cytotoxicity staining at end point should be saved in a subfolder “FLUO” without TP number.

(**B**) Illustration of the difference in segmentation of spheroid images when either ‘Invasion’ or ‘Growth’ strategy is employed by representative segmentation outputs of the two SImBA segmentation pipelines. Per row, left to right: the raw image, the segmentation mask when using the Invasion pipeline, and when using the Growth pipeline. The top two rows are brightfield images of spheroids suspended in medium (image sources see Table S4). Here the Growth strategy is employed as indicated in green box and this demonstrates how loose debris and small satellite spheroids are eliminated (LNCap, top row), as well as the microwell ring artefacts (SW837, row 2). The bottom two rows are phase-contrast invasion assay images of spheroids embedded in a collagen Type I ECM. Here the Invasion pipeline needs to be chosen (indicated in green) to also include the invading areas (single cells) for downstream invasion quantification. This figure illustrates how the choice in SImBA for either Growth or Invasion allows for a flexible and generic applicability across multiple different experimental set-ups.

(**C**) *Prescreen* quality control of SImBA segmentation. Only a representative subset of Hep3B spheroid *Prescreen* results is shown. Each figure panel displays the segmentation mask (black) with concentric, colour coded rings centred on the spheroid with increasing radius (1.5x-3x); these rings denote the selectable error-margin (EM) thresholds. Note that in an actual Prescreen a full montage of all spheroids is generated for every time point (t2 shown), enabling rapid checks for the entire experiment of (i) segmentation quality (e.g. blue square indicates over-segmentation of the ECM), (ii) ‘debris, artefacts’, to guide EM selection (red arrows, usually similarly present in all time points for a specific spheroid), and (iii) proximity of neighbouring spheroids (green arrows). EM-based thresholding implies that areas outside this radius multiplier are removed from further analysis, as also detailed in Supplementary Materials 1.1. Spheroid images failing quality control (QC) due to poor segmentation, excessive debris that is non-removable using EM-thresholding, or neighbouring spheroids in the image, can be excluded by the user before a *Prescreen* re-run (with same or different segmentation method) or before the full SImBA analysis. This incorporated SImBA segmentation-driven QC-step renders needless pre-evaluation of the image set by the user, but makes a fast exclusion of problematic images possible, now in a consistent (and due to EM thresholding, sufficiently subtle) manner based on specific issues or criteria after segmentation. It is aimed to meet the possibility to exclude problematic images but offers a way to do this based on segmentation outcome in a standardisable manner across users.

(**D**) Overview illustrating the diverse, high-quality visual output of SImBA. For all spheroids in the main directory SImBA generates 6 types of visual output: **(i)** three forms of manuscript-ready montages in time: raw, overlay of raw and segmented and binary view of segmented area, **(ii)** visualisation of the Invasion Mode Index (IMI) results for each timepoint (with outer bounding circle in black and, spheroid/cell-occupied area in white), **(iii)** visual comparison for each spheroid of growth or invasiveness in time based on an overlay of the total segmented area (spheroid and, if applicable, invaded cells) for the first ($t_0$, in red) and final time point ($t_{final}$, in green) **(iv)** (similar to (iii)) a composite image of spheroid outlines over time for each spheroids using colour code illustrating growth dynamics in a single view, **(v)** an overlay of the total segmented area of each spheroid on the raw image for each time point separately and **(vi)** protrusion images per timepoint based on skeletonization of the segmented spheroid overlayed on the raw spheroid image. The terminal voxels correspond to protrusion tips and their count approximates the number of protrusions. These visuals are for each spheroid in the experiment stored in the subfolders Montage, Index, MASK, Composite and Overlay, respectively as further clarified in Table S2. SImBA output overview. Additional visuals are stored in the Prescreen subfolder in the main directory.

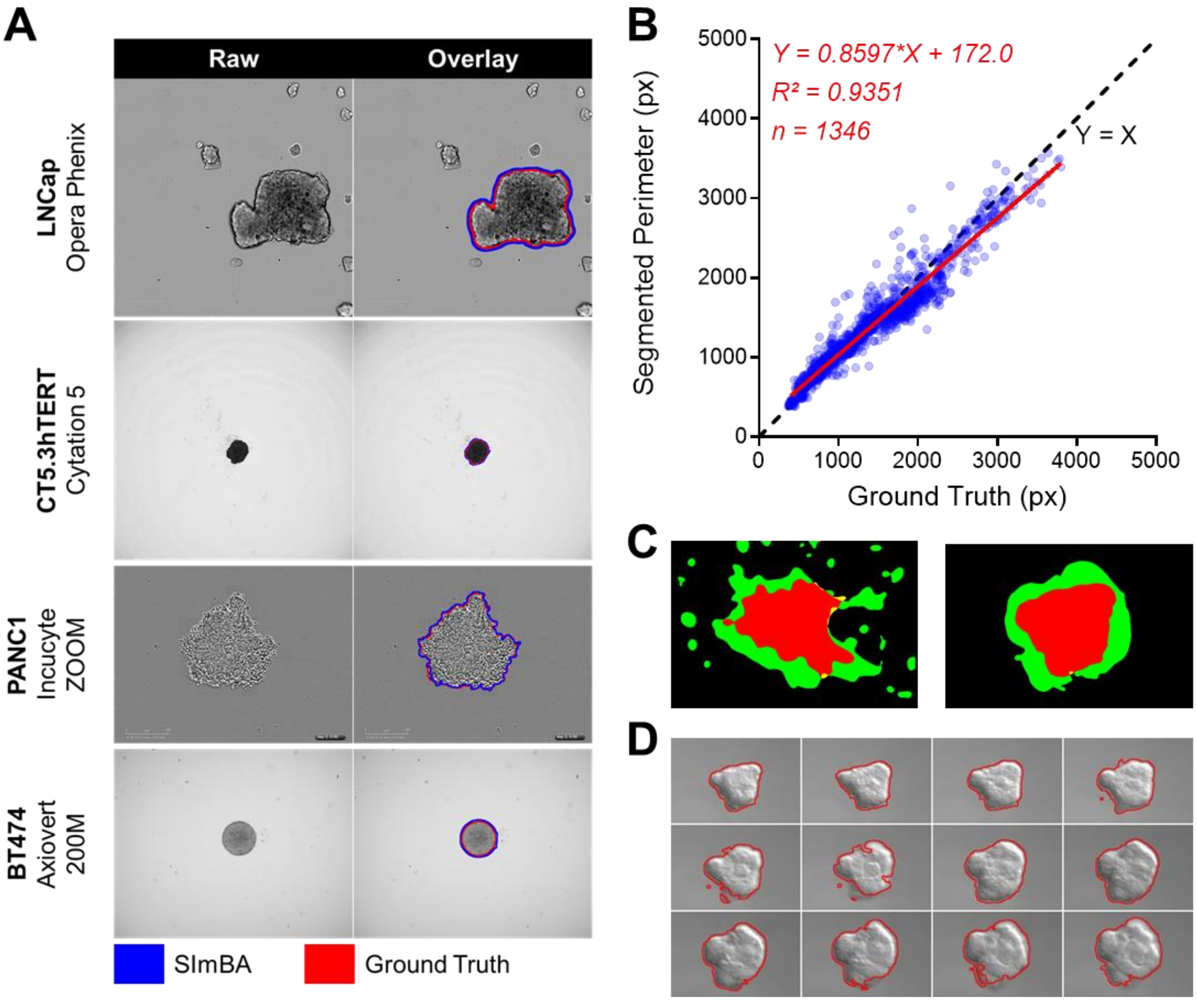

***Figure S2. SImBA produces accurate, ground-truth-comparable segmentations in spheroid and organoid 3D models.***

(**A**) Segmentation overlays on raw SLiMIA images show SImBA's automated segmentation contours (blue) alongside the SLiMIA ground-truth annotation contours (red), illustrating that SImBA produces highly accurate, ground-truth-comparable segmentations.

(**B**) Scatter plot of spheroid perimeter (px) quantified by SImBA versus SLiMIA manual ground truth displays strong correlation between manual and automated segmentation.

(**A,B**) see Table S4 for image sources used, and segmentation setting in SImBA.

(**C,D**) Murine mammary tumour organoid grown in Collagen type 1 (C, left) or Matrigel (C, right; D). Images are a selection of time points extracted from the movies CIL42156 and CIL42166 resp. (Cell Image Library, CIL (RRID:SCR_003510; https://doi.org/doi:10.7295/W9CIL42156) and segmented using SImBA (Find Edges, Sigma 6.5).

(**C**) SImBA visual output indicating the first timepoint red and the final timepoint in green to visually demonstrate the change in organoid morphology and invasive phenotype.

(**D**) SImBA visual output as a montage (overlay of the raw image with the segmented area) This illustrates that next to spheroids, also organoids can be processed by SImBA-SiQuAl, largely expanding the application range.

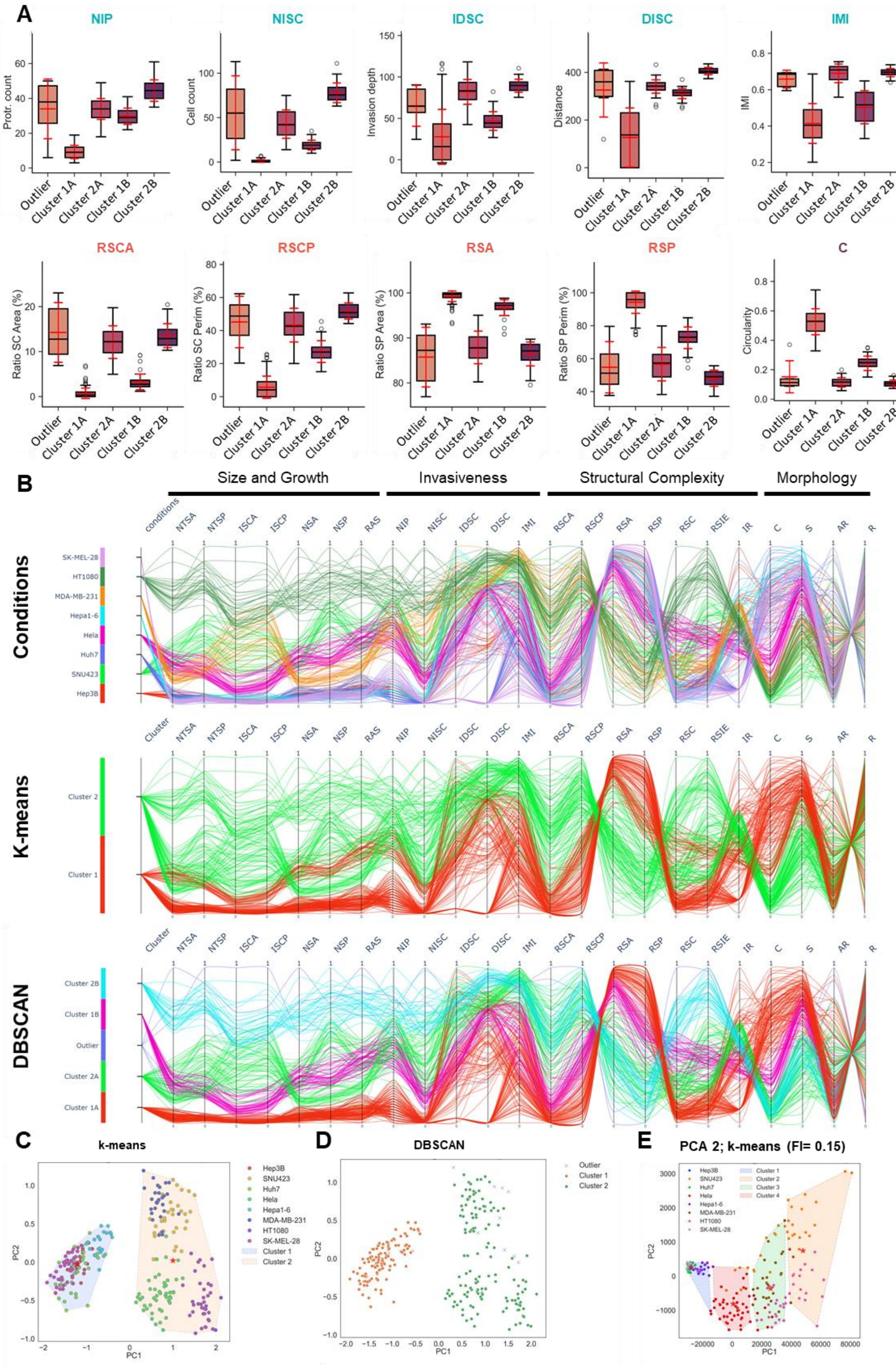
A
NIP
Protr. count
NISC
Cell count
IDSC
Invasion depth
DISC
Distance
IMI
IMI
RSCA
Ratio SC Area (%)
RSCP
Ratio SC Perim (%)
RSA
Ratio SP Area (%)
RSP
Ratio SP Perim (%)
C
Circularity
Outlier
Cluster 1A
Cluster 2A
Cluster 1B
Cluster 2B
B
Size and Growth
Invasiveness
Structural Complexity
Morphology
Conditions
conditions
NTSA
NTSP
ISCA
ISCP
NSA
NSP
RAS
NIP
NISC
IDSC
DISC
IMI
RSCA
RSCP
RSA
RSP
RSC
RSIE
IR
C
S
AR
R
SK-MEL-28
HT1080
MDA-MB-231
Hepa1-6
Hela
Huh7
SNU423
Hep3B
K-means
Cluster
Cluster 2
Cluster 1
DBSCAN
Cluster 2B
Cluster 1B
Outlier
Cluster 2A
Cluster 1A
C
k-means
PC1
PC2
Cluster 1
Cluster 2
D
DBSCAN
Outlier
Cluster 1
Cluster 2
E
PCA 2; k-means (FI= 0.15)
Cluster 3
Cluster 4

***Figure S3. Case study 1: SImBA–SiQuAl identifies distinct spheroid invasion phenotypes***

(**A**) DBSCAN cluster boxplots of key Feature Importance (FI) ranked features, including the distance of invasion and invasion depth of single cells (DISC, IDSC resp.) and the invasion mode index (IMI), the number of invading single cells (NISC), protrusions (NIP), the ratios of single-cell/spheroid to total area and perimeter (RSCA, RSCP, RSA and RSP), and circularity (C) at 24h ($t_1$).

(**B**) Parallel-coordinate plots of min-max scaled SiQuAl features, grouped by feature category and coloured by condition, k-means cluster or DBSCAN cluster. Condition-based visualization shows strong cell line-specific phenotypic differences, while k-means separates the dataset into two broad phenotypic groups. DBSCAN further resolves these into more detailed sub-clusters.

(**D, E**) PCA with clustering (k-means (D), DBSCAN (E)) separates cell lines into two major groups after 48h ($t_2$).

(**F**) FI filtered (threshold 0.15) PCA with k-means clustering partly resolves the loss of cluster resolution at 48h.

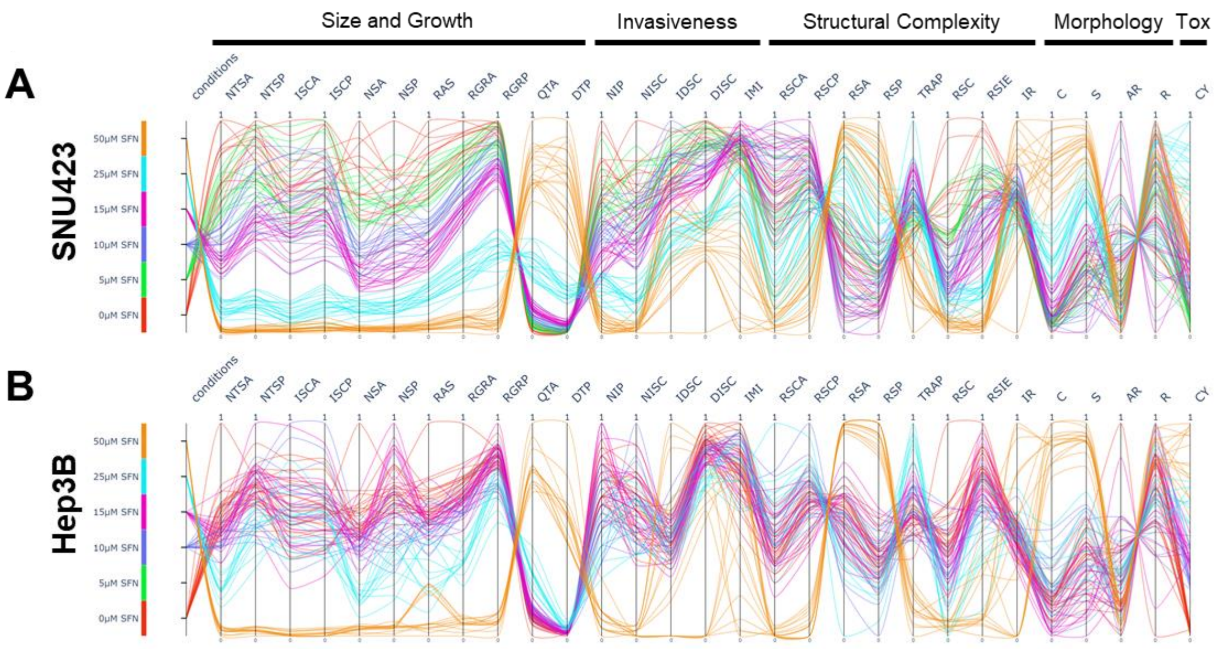

***Figure S4. Case study 2: Capturing Distinct Sorafenib-Induced Phenotypic Responses in HCC Cell Lines***

Parallel-coordinate plots of min-max-scaled SiQuAl features for SFN-treated Hep3B (**A**) and SNU423 (**B**) spheroids, coloured by condition. These plots visualize the dose-dependent phenotypic response to SFN. SNU423 spheroids show an overall dose-dependent phenotypic shift across most SiQuAl parameters. In contrast, for Hep3B spheroids, the low-dose SFN effect is limited to a decrease in size- and growth-related features only at 25µM while invasion features display little to no dose-dependency. 50µM SFN induces maximal toxicity in both cell lines. (Tox = Cytotoxicity)

### SUPPLEMENTARY TABLES

***Table S1. Features (parameters) extracted by SImBA and structured/normalised by SiQuAl.***

Features are provided with a description and, where appropriate, the corresponding formula/mathematical definition, unit and feature category (Cat). Features can be grouped into 5 categories including: Size and Growth (SG), Invasiveness (I), Structural complexity (SC), Morphology (M), and cytotoxicity (C).

| Feature name | Description | Mathematical definition | Unit | Cat |
|---|---|---|---|---|
| **Total Spheroid Area (TSA)** | This is the non-normalised summed area of the spheroid and single cells as measured by SImBA. | | px² | SG |
| **Normalized Total Spheroid Area (NTSA)** | This is the summed area of the spheroid and single cells at time point n, normalized to the *TSA* of the first timepoint ($t_0$). Normalization is performed by SiQuAl. | $NTSA_{tn} = \frac{TSA_{tn}}{TSA_{t0}} \times 100$ | % | SG |
| **Spheroid Area (SA)** | Area of the main spheroid mass without single cells, measured on images in which single cells were graphically removed. This is the non-normalised area as measured by SImBA. | $SA = TSA - ISCA$ | px² | SG |
| **Normalized Spheroid Area (NSA)** | Area of the spheroid without single cells (*SA*) at time point n normalized to the *SA* of the first timepoint ($t_0$). Normalization is performed by SiQuAl. | $NSA_{tn} = \frac{SA_{tn}}{SA_{t0}} \times 100$ | % | SG |
| **Invading Single Cell Area (ISCA)** | Area of all invading single cells, measured by SImBA on images in which the main spheroid mass was graphically removed. | $ISCA = TSA - SA$ | px² | SG |
| **Total Spheroid Perimeter (TSP)** | This is the summed perimeter of the area occupied by the main spheroid mass and invading single cells as measured by SImBA. | | px | SG |
| **Normalized Total Spheroid Perimeter (NTSP)** | This is the perimeter of area occupied by the spheroid and single cells *(TSP)* at timepoint n ($t_n$) normalized to the *TSP* of the first timepoint ($t_0$). Normalization is performed by SiQuAl. | $NTSP_{tn} = \frac{TSP_{tn}}{TSP_{t0}} \times 100$ | % | SG |
| **Spheroid Perimeter (SP)** | Perimeter of the main spheroid mass, measured on images in which the single cells were graphically removed. This is the non-normalised perimeter as measured by SImBA. | $SP = TSP - ISCP$ | px | SG |
| **Normalized Spheroid Perimeter (NSP)** | Perimeter of the main spheroid mass *(SP)* at timepoint n normalized to *SP* of | $NSP_{tn} = \frac{SP_{tn}}{SP_{t0}} \times 100$ | % | SG |

| | | | | |
|---|---|---|---|---|
| | the first timepoint ($t_0$). Normalization is performed by SiQuAl. | | | |
| **Invading Single Cell Perimeter (ISCP)** | Summed perimeter of all invading single cells, measured by SImBA on images in which the spheroid was graphically removed. | $ISCP = TSP - SP$ | px | SG |
| **Ratio of Single Cell Area over Total Area (RSCA)** | Ratio of the *ISCA* over the *TSA* as a measure of single cell invasiveness. | $RSCA_{tn} = \frac{ISCA_{tn}}{TSA_{tn}}$ | | SC |
| **Ratio of Single Cell Perimeter over Total Perimeter (RSCP)** | Ratio of the *ISCP* over the *TSP* as a measure of single cell invasiveness. | $RSCP_{tn} = \frac{ISCP_{tn}}{TSP_{tn}}$ | | SC |
| **Ratio of Spheroid Area over Total Area (RSA)** | Ratio of the *SA* over the *TSA.* | $RSA_{tn} = \frac{SA_{tn}}{TSA_{tn}}$ | | SC |
| **Ratio of Spheroid Perimeter over Total Perimeter (RSP)** | Ratio of the *SP* over the *TSP.* | $RSP_{tn} = \frac{SP_{tn}}{TSP_{tn}}$ | | SC |
| **Radius of the averaged spheroid (RAS)** | The radius of the "averaged spheroid", measured but fitting the average spheroid. This is a fitted circle with its centre in the centre of mass of the spheroid and an area equal to the SA. | | px | SG |
| **Radius of the spheroid's core (RSC)** | The radius of the maximum inscribed circle within the spheroid | | px | SC |
| **Radius of the spheroid's invasive edge (RSIE)** | The radius of the minimum outer bounding circle (OBC) that inscribes all areas in the image | | px | SC |
| **Number of invading single cells (NISC)** | The number of areas invading the matrix excluding the spheroid after watershed filter. | | | I |
| **Number of invading protrusions (NIP)** | The number of protrusions that emerge from the spheroid's surface using the end voxels of the FIJI Skeletonize plugin (*legacy:ij.plugin.filter.Binary("skel")*) | | | I |
| **Circularity (C)** | A value of 1 indicates a perfect circle. The closer to 0, the less circular the spheroid becomes. Highly invasive spheroids have a low circularity due to irregular, i.e. larger perimeter. | $C = 4\pi \times \frac{SA}{TSP^2}$ | | M |

| | | | | |
|---|---|---|---|---|
| **Solidity (S)** | A measure of the density of the spheroid; used to describe the irregularity of the spheroid's perimeter. Low value for invasive spheroids with strongly irregular perimeter | $S = \frac{SA}{Convex\ SA}$ | | M |
| **Aspect Ratio (AR)** | A measure of elongation or directional expansion/invasion of the spheroid. | $AR = \frac{\text{major axis}}{\text{minor axis}}$ | | M |
| **Roundness (R)** | A measure of elongation or directional expansion/invasion of the spheroid. | $R = 4 \times \frac{SA}{\pi\ \times \text{major axis}^2} = \frac{1}{AR}$ | | M |
| **Distance of invasion of single cells (DISC)** | The mean of the distances (d) between the centroid of the spheroid and the centroid of invading areas. | $\text{DISC} = \frac{1}{NISC}\sum_{i=1}^{d_n} d_i$ | px | I |
| **Invasion Depth of single cells (IDSC)** | The mean of the distances (db) between the centroid of the invading areas and the boundary of the 'averaged spheroid' (as defined above for RAS). | $\text{IDSC} = \frac{1}{NISC}\sum_{i=1}^{db_n} db_i$ | px | I |
| **Invasion Ratio (IR)** | The ratio of IDSC over DISC as a measure for invasion normalized to spheroid area | $IR = \frac{IDSC}{DISC}$ | | SC |
| **Invasion Mode Index (IMI)** | The ratio of the cell free area within the outer bounding circle (OBC) over the total area within the OBC as an indication of invasion mode as described in Supplementary Figure 4C | $IMI = \frac{Cell\ free\ area}{Total\ area\ in\ the\ OBC}$ | | I |
| **Relative Growth Rate of the Area (RGRA)** | A standardized, unbiased measure of growth of the TSA (summed area of spheroid and invading areas) excluding inherent differences in scale. This parameter is useful to compare between experiments/conditions Timepoint n can be altered by the user, but should be default the final timepoint ($t_n$). | $RGR_A = \frac{\ln TSA_n - \ln TSA_0}{t_n - t_0}$ | $px^2/h$ | SG |
| **Relative Growth Rate of the Perimeter (RGRP)** | A standardized, unbiased measure of growth of the perimeter TSP (perimeter of summed area occupied by spheroid and invading areas) without inherent differences in scale. Timepoint n can be altered by the user, but should be default the final timepoint ($t_n$). | $RGR_P = \frac{\ln TSP_n - \ln TSP_0}{t_n - t_0}$ | px/h | SG |
| **Doubling Time of the Perimeter (DTP)** | The $DT_P$ describes the (extrapolated) time needed for the perimeter (TSP) of the spheroid to double. | $DT_P = \frac{\ln 2}{RGR_p}$ | h | SG |

| | | | | |
|---|---|---|---|---|
| **Quadrupling Time of the Area (QTA)** | The $QT_A$ describes the (extrapolated) time needed to increase the area (TSA) four times. | $QT_A = \frac{2 * \ln 2}{RGR_A}$ | h | SG |
| **Time dependent Ratio of Area over Perimeter (TRAP)** | The ratio of the quadrupling time of the area over the doubling time of the perimeter as a time dependent indication of invasion mode. For perfect circular growth TRAP = 1 | $TRAP = \frac{QT_A}{DT_P}$ | | SC |
| **Cytotoxicity (CY)** | Describes the mean intensity of the fluorescent (cytotoxicity) dye within the TSA of the spheroid. This is the sum of the gray values (GV) of all n pixels ($P_n$) in the area divided by the number of pixels (np). | $CY = \frac{\sum_{P_0}^{P_n} GV}{\sum_1^n np}$ | | C |

***Table S2. SImBA output overview***
Comprehensive overview of all output files (numerical and visual) generated by SImBA.
**File 1-**8: These files are numerical data files stored in a separate *Measurements* folder in the main directory. These files contain numerical output for the named features, and are organized in such a way that timepoints of each spheroid from the main directory are in consecutive rows.
**File 9-13:** These files are the results of the final Prescreen run, stored in a separate *Prescreen_Results* folder in the main directory.
**File 14-75:** These files describe all output generated by SImBA and stored per spheroid (in the spheroid subfolder) for an example image (here QA01_01) imaged over 5 timepoints (t0, t1, t2, t3, t4)
Abbreviations: FN: File number, SF: subfolder, IO: Intermediate Output, FO: Final Output

| | File | Folder\SF | Description | Raw/ IO/ FO/ SF | Example Image |
|---|---|---|---|---|---|
| **1** | SImBA_log_yyyy-mm-dd_hh-mm-ss.txt | Main_Directory\ Measurements | Contains all log statements that were printed in the log window during a SImBA run | FO | text |
| **1** | Results_ segmentation .csv | Main_Directory\ Measurements | Contains raw numerical data of the following features: Area Total, Area Spheroid, Area Single Cells, Perimeter Total, Perimeter Spheroid, Perimeter Single Cells, Ratio Single Cell Area (%), Ratio Spheroid Area (%), Ratio Single Cell Perimeter (%), Ratio Spheroid Perimeter (%), Circularity, Solidity, Roundness, Aspect Ratio for each spheroid for all timepoints in subsequent rows. The different features are sorted in subsequent columns. | FO | Combined Numerical Output |
| **2** | Radius_Mean_ Core_ Invasion.csv | Main_Directory\ Measurements | Contains raw numerical data of the following features: Mean radius, Core radius, Invasive radius for each spheroid for all timepoints in subsequent rows. The different features are sorted in subsequent columns. | FO | Combined Numerical Output |
| **3** | TRAP_ Results.csv | Main_Directory\ Measurements | Contains raw data of the following features: Relative Growth Rate Area, Relative Growth Rate Perimeter, Doubling Time Perimeter, Quadrupling Time Area, TRAP at the timepoint chosen to assess growth kinetics (final timepoint) for each spheroid in | FO | Combined Numerical Output |

| | | | | | |
|---|---|---|---|---|---|
| | | | subsequent rows. The different features are sorted in subsequent columns. | | |
| **4** | Cell_count.csv | Main_Directory\ Measurements | Contains raw Cell count data for each spheroid for all timepoints in subsequent rows. | FO | Combined Numerical Output |
| **5** | Invasion_ Depth.csv | Main_Directory\ Measurements | Contains raw data of the following features: Distance, Invasion depth, Invasion Ratio for each spheroid for all timepoints in subsequent rows. The different features are sorted in subsequent columns. | FO | Combined Numerical Output |
| **6** | Invasion_Mode_ Index.csv | Main_Directory\ Measurements | Contains raw IMI data for each spheroid for all timepoints in subsequent rows. The different features are sorted in subsequent columns. | FO | Combined Numerical Output |
| **7** | Protrusion_ count.csv | Main_Directory\ Measurements | Contains raw Protrusion count data for each spheroid for all timepoints in subsequent rows. The different features are sorted in subsequent columns. | FO | Combined Numerical Output |
| **8** | Results Cytotoxicity.csv | Main_Directory\ Measurements | Contains raw Cytoxicity data | FO | Combined Numerical Output |
| **9** | montage t0.jpg | Main_Directory\ Prescreen_ Results | Montage of the segmentations of all spheroids at t0. On top of the spheroids are concentric circles overlayed as guide for choosing an appropriate Error Margin. | FO | 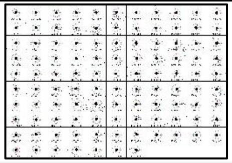 |
| **10** | montage t1.jpg | Main_Directory\ Prescreen_ Results | Montage of the segmentations of all spheroids at t1. On top of the spheroids are concentric circles overlayed as guide for choosing an appropriate Error Margin. | FO | 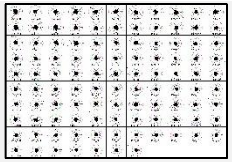 |
| **11** | montage t2.jpg | Main_Directory\ Prescreen_ Results | Montage of the segmentations of all spheroids at t2. On top of the spheroids are concentric circles overlayed as guide for choosing an appropriate Error Margin. | FO | 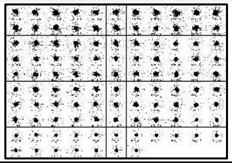 |

| | | | | | |
|---|---|---|---|---|---|
| **12** | montage t3.jpg | Main_Directory\ Prescreen_ Results | Montage of the segmentations of all spheroids at t3. On top of the spheroids are concentric circles overlayed as guide for choosing an appropriate Error Margin. | FO | 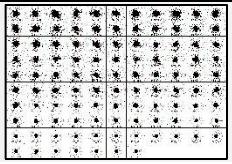 |
| **13** | montage t4.jpg | Main_Directory\ Prescreen_ Results | Montage of the segmentations of all spheroids at t4. On top of the spheroids are concentric circles overlayed as guide for choosing an appropriate Error Margin. | FO | 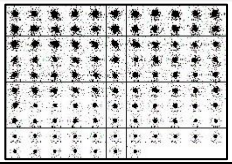 |
| **14** | QA01_01 t0.tif | Main_Directory\ QA01_01 | Raw image of Spheroid QA01_01 at timepoint t0 | RAW | 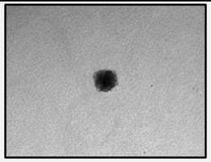 |
| **15** | QA01_01 t1.tif | Main_Directory\ QA01_01 | Raw image of Spheroid QA01_01 at timepoint t1 | RAW | 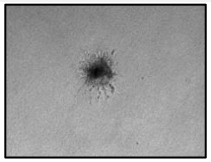 |
| **16** | QA01_01 t2.tif | Main_Directory\ QA01_01 | Raw image of Spheroid QA01_01 at timepoint t2 | RAW | 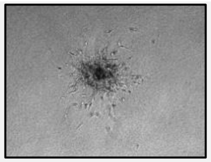 |
| **17** | QA01_01 t3.tif | Main_Directory\ QA01_01 | Raw image of Spheroid QA01_01 at timepoint t3 | RAW | 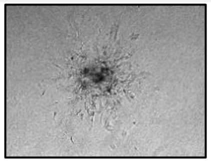 |
| **18** | QA01_01 t4.tif | Main_Directory\ QA01_01 | Raw image of Spheroid QA01_01 at timepoint t4 | RAW | 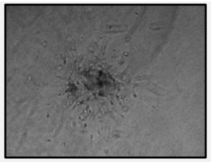 |
| **19** | QA01_01\FLUO\ | Main_Directory\ QA01_01 | Original (optional) subfolder. This subfolder contains raw fluorescent cytotoxicity staining at the final timepoint | SF | Subfolder |

| | | | | | |
|---|---|---|---|---|---|
| **20** | QA01_01\aaa_ Segmentation\ | Main_Directory\ QA01_01 | Newly created subfolder. This subfolder contains the original segmentation masks before any adjustments are made. | SF | Subfolder |
| **21** | QA01_01\Adj_ Segmentation\ | Main_Directory\ QA01_01 | Newly created subfolder. This subfolder contains adjusted new segmentation masks. Adjusted means that areas outside of the Error Margin are removed. | SF | Subfolder |
| **22** | QA01_01\ Composite\ | Main_Directory\ QA01_01 | Newly created subfolder. This subfolder contains the composite of the outline of the segmentation masks. | SF | Subfolder |
| **23** | QA01_01\ Index\ | Main_Directory\ QA01_01 | Newly created subfolder. This subfolder contains the images created to measure the invasion mode index | SF | Subfolder |
| **24** | QA01_01\ MASK\ | Main_Directory\ QA01_01 | Newly created subfolder. This subfolder contains the t0 versus tfinal mask. | SF | Subfolder |
| **25** | QA01_01\ Montage\ | Main_Directory\ QA01_01 | Newly created subfolder. This subfolder contains montages of raw and segmented images as well as their overlay. | SF | Subfolder |
| **26** | QA01_01\New_ Segmentation\ | Main_Directory\ QA01_01 | Newly created subfolder. This subfolder contains the original but doubled segmentation masks. | SF | Subfolder |
| **27** | QA01_01\ Overlay\ | Main_Directory\ QA01_01 | Newly created subfolder. This subfolder contains overlay images of segmentation masks and protrusion mask. | SF | Subfolder |
| **28** | QA01_01\ROI\ | Main_Directory\ QA01_01 | Newly created subfolder. This subfolder contains the raw segmented region of interest data (as stored from the ROI manager in FIJI) | SF | Subfolder |
| **29** | QA01_01.tif | Main_Directory\ QA01_01\FLUO | Raw fluorescent image of the cytotoxicity staining of Spheroid QA01_01 at timepoint t4 (t Final) | RAW | 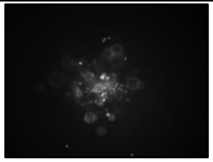 |

| | | | | | |
|---|---|---|---|---|---|
| **30** | t0 Seg.tif | Main_Directory\QA01_01\aaa_Segmentation | Original Segmentation mask of spheroid QA01_01 at timepoint t0 | IO and FO | 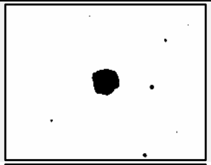 |
| **31** | t1 Seg.tif | Main_Directory\QA01_01\aaa_Segmentation | Original Segmentation mask of spheroid QA01_01 at timepoint t1 | IO and FO | 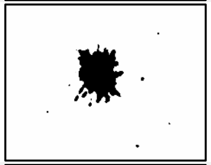 |
| **32** | t2 Seg.tif | Main_Directory\QA01_01\aaa_Segmentation | Original Segmentation mask of spheroid QA01_01 at timepoint t2 | IO and FO | 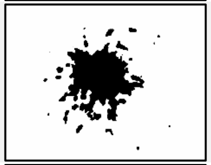 |
| **33** | t3 Seg.tif | Main_Directory\QA01_01\aaa_Segmentation | Original Segmentation mask of spheroid QA01_01 at timepoint t3 | IO and FO | 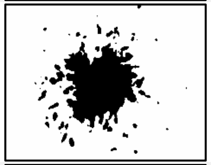 |
| **34** | t4 Seg.tif | Main_Directory\QA01_01\aaa_Segmentation | Original Segmentation mask of spheroid QA01_01 at timepoint t4 | IO and FO | 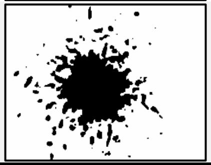 |
| **35** | t0 newSeg.tif | Main_Directory\QA01_01\New_Segmentation | New Segmentation mask of spheroid QA01_01 at timepoint t0. This new segmentation mask is an intermediate output as it contains exactly the same segmentation as in the original mask, however, the boundries of the images are extended to double the image size. This doubling is to prevent an underestimation of the Invasion Mode Index. As the IMI relies on drawing the outer bounding circle around the spheroid and its single cells, this circle often falls outside the original image boundaries for large/growing spheroids. | IO | 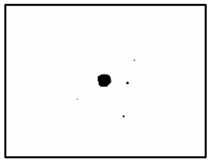 |

| | | | | | |
|---|---|---|---|---|---|
| **36** | t1 newSeg.tif | Main_Directory \QA01_01\New_ Segmentation | New Segmentation mask of spheroid QA01_01 at timepoint t1 | IO | 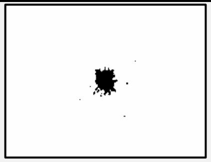 |
| **37** | t2 newSeg.tif | Main_Directory \QA01_01\New_ Segmentation | New Segmentation mask of spheroid QA01_01 at timepoint t2 | IO | 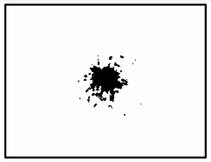 |
| **38** | t3 newSeg.tif | Main_Directory \QA01_01\New_ Segmentation | New Segmentation mask of spheroid QA01_01 at timepoint t3 | IO | 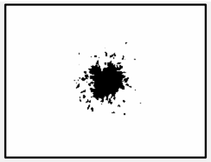 |
| **39** | t4 newSeg.tif | Main_Directory \QA01_01\New_ Segmentation | New Segmentation mask of spheroid QA01_01 at timepoint t4 | IO | 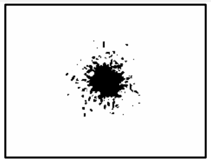 |
| **40** | t0 AdjustedSeg.tif | Main_Directory\ QA01_01\Adj_ Segmentation | Adjusted Segmentation mask of spheroid QA01_01 at timepoint t0. Using the user provided Error Margin (EM), Areas/debris that falls outside of the radius * EM is removed. This adjusted segmentation takes the new segmentation (double image size, same spheroid size) as input. | IO | 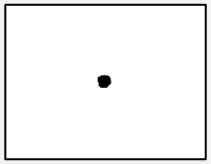 |
| **41** | t1 AdjustedSeg.tif | Main_Directory\ QA01_01\Adj_ Segmentation | Adjusted Segmentation mask of spheroid QA01_01 at timepoint t1. | IO | 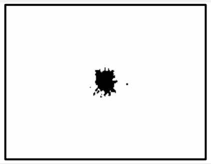 |
| **42** | t2 AdjustedSeg.tif | Main_Directory\ QA01_01\Adj_ Segmentation | Adjusted Segmentation mask of spheroid QA01_01 at timepoint t2. | IO | 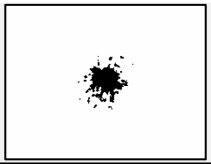 |

| | | | | | |
|---|---|---|---|---|---|
| **43** | t3 AdjustedSeg.tif | Main_Directory\ QA01_01\Adj_ Segmentation | Adjusted Segmentation mask of spheroid QA01_01 at timepoint t3. | IO | 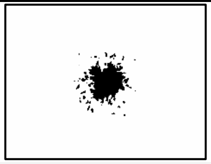 |
| **44** | t4 AdjustedSeg.tif | Main_Directory\ QA01_01\Adj_ Segmentation | Adjusted Segmentation mask of spheroid QA01_01 at timepoint t4. | IO | 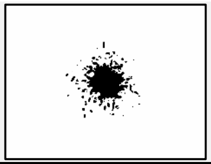 |
| **45** | QA01_01 Composite Outlines.png | Main_Directory\ QA01_01\ Composite | Composite image (.png) of the Masks/Segmentation outlines of all timepoints (here 5 time points: t0-t4) in 1 image. This composite image can contain up to 7 different outlines, for time series with more than 7 timepoints, use the "t0_versus_tfinal.tif". (t0: red, t1: green, t2: blue, t3: cyan, t4: white) | FO | 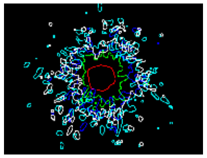 |
| **46** | QA01_01 Composite Outlines.tif | Main_Directory\ QA01_01\ Composite | Composite image (.tif) of the Masks/Segmentation outlines of all timepoints (here 5 time points: t0-t4) in 1 image. This composite image can contain up to 7 different outlines, for time series with more than 7 timepoints, use the "t0_versus_tfinal.tif". (t0: red, t1: green, t2: blue, t3: cyan, t4: white) | FO | 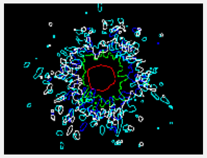 |
| **47** | t0_cellFree.tif | Main_Directory\ QA01_01\Index | Invasion Mode Index figures. Mainly used as intermediate output to calcultate the ratio of cell free area over total area, but can be used as final output to visualize the extend of invasion. Both cell free and full circle images are based on the adjusted segmentation masks: fullCircle.tif images contain the filled in outer bounding circle of the spheroid and its invading cells. Similarly, cellFree.tif contain the same area with the spheroid and single cell areas substracted. | IO and FO | 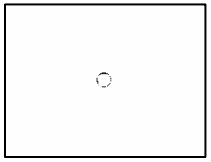 |

| | | | | | |
|---|---|---|---|---|---|
| | | | Cell free area within the outer bounding circle of spheroid QA01_01 at timepoint t0. | | |
| **48** | t0_fullCircle.tif | Main_Directory\QA01_01\Index | Total filled in area within the outer bounding circle of spheroid QA01_01 at timepoint t0. | IO and FO | 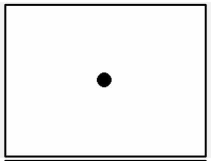 |
| **49** | t1_cellFree.tif | Main_Directory\QA01_01\Index | Cell free area within the outer bounding circle of spheroid QA01_01 at timepoint t1. | IO and FO | 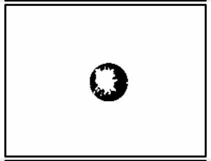 |
| **50** | t1_fullCircle.tif | Main_Directory\QA01_01\Index | Total filled in area within the outer bounding circle of spheroid QA01_01 at timepoint t1. | IO and FO | 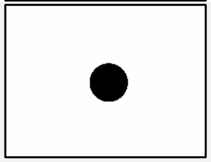 |
| **51** | t2_cellFree.tif | Main_Directory\QA01_01\Index | Cell free area within the outer bounding circle of spheroid QA01_01 at timepoint t2. | IO and FO | 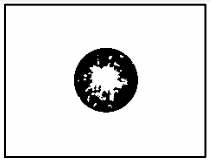 |
| **52** | t2_fullCircle.tif | Main_Directory\QA01_01\Index | Total filled in area within the outer bounding circle of spheroid QA01_01 at timepoint t2. | IO and FO | 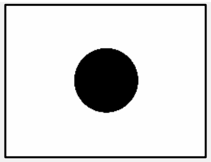 |
| **53** | t3_cellFree.tif | Main_Directory\QA01_01\Index | Cell free area within the outer bounding circle of spheroid QA01_01 at timepoint t3. | IO and FO | 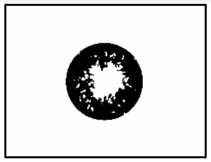 |
| **54** | t3_fullCircle.tif | Main_Directory\QA01_01\Index | Total filled in area within the outer bounding circle of spheroid QA01_01 at timepoint t3. | IO and FO | 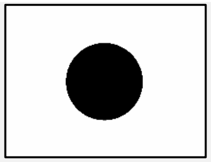 |

| | | | | | |
|---|---|---|---|---|---|
| **55** | t4_cellFree.tif | Main_Directory\QA01_01\Index | Cell free area within the outer bounding circle of spheroid QA01_01 at timepoint t4. | IO and FO | 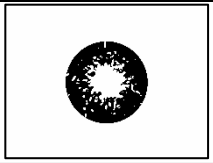 |
| **56** | t4_fullCircle.tif | Main_Directory\QA01_01\Index | Total filled in area within the outer bounding circle of spheroid QA01_01 at timepoint t4. | IO and FO | 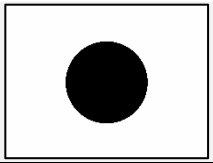 |
| **57** | t0_versus_ tfinal.tif | Main_Directory\QA01_01\MASK | Composite image (.tif) of the Masks of the first and last timepoints (here t0 and t4) in 1 image. This composite image is a visual representation of spheroid change and can be used as simplified "QA01_01 Composite Outlines". for time series that include more than 7 timepoints use this instead of "QA01_01 Composite Outlines". (t0: red, t'/tfinal: green) | FO | 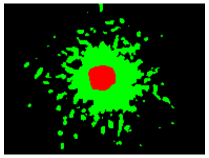 |
| **58** | QA01_01 Montage Raw data.jpg | Main_Directory\QA01_01\Montage | Montage (.jpg) of the raw images for each timepoint. | FO | 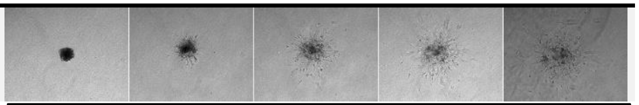 |
| **59** | QA01_01 Montage Segmented data.jpg | Main_Directory\QA01_01\Montage | Montage (.jpg) of the segmentation masks for each timepoint. | FO | 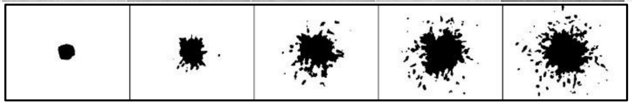 |
| **60** | QA01_01 Montage Overlay data.jpg | Main_Directory\QA01_01\Montage | Montage (.jpg) of the segmentation overlay images for each timepoint. | FO | 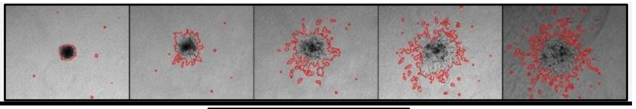 |
| **61** | QA01_01 t0.jpg | Main_Directory\QA01_01\Overlay | Raw images of spheroid QA01_01 at timepoint t0 overlayed with the appropriate segmentation mask in red | FO | 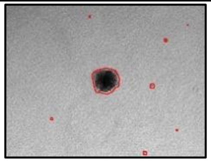 |

| | | | | | |
|---|---|---|---|---|---|
| **62** | QA01_01 t1.jpg | Main_Directory\ QA01_01\Overlay | Raw image of spheroid QA01_01 at timepoint t1 overlayed with the appropriate segmentation mask in red | FO | 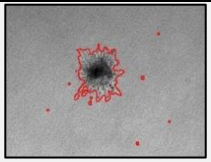 |
| **63** | QA01_01 t2.jpg | Main_Directory\ QA01_01\Overlay | Raw image of spheroid QA01_01 at timepoint t2 overlayed with the appropriate segmentation mask in red | FO | 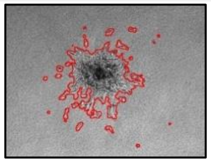 |
| **64** | QA01_01 t3.jpg | Main_Directory\ QA01_01\Overlay | Raw image of spheroid QA01_01 at timepoint t3 overlayed with the appropriate segmentation mask in red | FO | 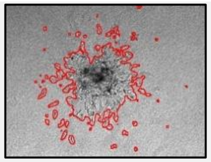 |
| **65** | QA01_01 t4.jpg | Main_Directory\ QA01_01\Overlay | Raw image of spheroid QA01_01 at timepoint t4 overlayed with the appropriate segmentation mask in red | FO | 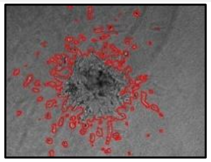 |
| **66** | t0 protr.jpg | Main_Directory\ QA01_01\Overlay | Raw image of spheroid QA01_01 at timepoint t0 overlayed with the appropriate protrusion (skeletonize-) mask in red | FO | 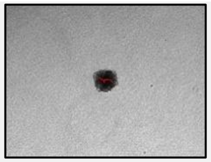 |
| **67** | t1 protr.jpg | Main_Directory\ QA01_01\Overlay | Raw image of spheroid QA01_01 at timepoint t1 overlayed with the appropriate protrusion (skeletonize-) mask in red | FO | 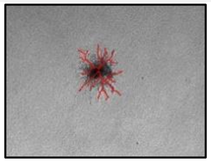 |
| **68** | t2 protr.jpg | Main_Directory\ QA01_01\Overlay | Raw image of spheroid QA01_01 at timepoint t2 overlayed with the appropriate protrusion (skeletonize-) mask in red | FO | 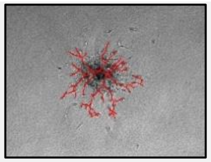 |

| | | | | | |
|---|---|---|---|---|---|
| **69** | t3 protr.jpg | Main_Directory\ QA01_01\Overlay | Raw image of spheroid QA01_01 at timepoint t3 overlayed with the appropriate protrusion (skeletonize-) mask in red | FO | 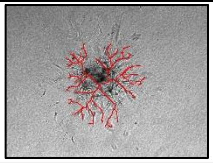 |
| **70** | t4 protr.jpg | Main_Directory\ QA01_01\Overlay | Raw image of spheroid QA01_01 at timepoint t4 overlayed with the appropriate protrusion (skeletonize-) mask in red | FO | 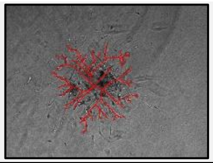 |
| **71** | t0 ROI.zip | Main_Directory\ QA01_01\ROI | Segmentation ROI of spheroid QA01_01 at timepoint t0. These ROIs are stored to perform all downstream measurements and mask transformations | IO | \ |
| **72** | t1 ROI.zip | Main_Directory\ QA01_01\ROI | Segmentation ROI of spheroid QA01_01 at timepoint t1. | IO | \ |
| **73** | t2 ROI.zip | Main_Directory\ QA01_01\ROI | Segmentation ROI of spheroid QA01_01 at timepoint t2. | IO | \ |
| **74** | t3 ROI.zip | Main_Directory\ QA01_01\ROI | Segmentation ROI of spheroid QA01_01 at timepoint t3. | IO | \ |
| **75** | t4 ROI.zip | Main_Directory\ QA01_01\ROI | Segmentation ROI of spheroid QA01_01 at timepoint t4. | IO | \ |

*Table S3. SiQuAl output overview*

Comprehensive overview of the files and subfolders generated by SiQuAl. For each file or file group, the table indicates its location within the directory structure, a brief description of its contents, and whether it represents intermediate output (IO), final output (FO), or subfolder (SF).

| File(s)/folder(s) | Folder\SF | Description | IO/ FO/ SF |
|---|---|---|---|
| **SiQuAl_log.txt** | Measurements (Main Folder) | This text file contains time stamped comments from the SiQuAl analysis. It prints the individual steps executed and provides the ranked FI-lists of features above the threshold. | FO |
| **SiQuAl_error_log.txt** | Measurements (Main Folder) | This text file contains time stamped error messages of errors that occurred while running SiQuAl. | FO |
| **areaSinglCells t0.csv; areaSinglCells t1.csv; areaSinglCells t2.csv; circularity t0.csv; circularity t1.csv; circularity t2.csv; …** | Output | .csv files with structured (and normalized) numerical output per parameter per timepoint | IO |
| **Output\Original** | Output | SF | SF |
| **Output\xlsx** | Output | SF | SF |
| **Results_segmentation.xlsx, TRAP_Results.xlsx, Cell_count.xlsx, Invasion_Depth.xlsx, Invasion_Mode_Index.xlsx, Protrusion_count.xlsx, Radius_Mean_Core_Invasion.xlsx** | Output\Original | .xlsx files with the original, Non-normalized structured numerical output .xlsx files: columns contain condition per timepoint (e.g. Hep3B t0; Hep3B t1; Hep3B t2) per feature. Individual parameters are sorted in separate xlsx sheets. | IO |
| **areaTotal.xlsx, aspectRatio.xlsx, Cell_count.xlsx, circularity.xlsx, areaSingleCells.xlsx, areaSpheroid.xlsx, …** | Output\xlsx | .xlsx files with normalized (where required) numerical output per individual feature. Columns contain condition per timepoint (e.g. Hep3B t0; Hep3B t1; Hep3B t2). Individual parameters are sorted in separate xlsx files. | IO |

| | | | |
|---|---|---|---|
| **Abs_Feature_importance_TP0.xlsx, Abs_Feature_importance_TP1.xlsx, Abs_Feature_importance_TP2.xlsx, …** | PCA (for k-means and DBSCAN analysis) | Absolute PC loadings for al principal components for each feature represented as a matrix in an .xlsx file | FO and IO |
| **Abs_Feature_importance_TP0.png, Abs_Feature_importance_TP1.png, Abs_Feature_importance_TP2.png, …** | PCA (for k-means and DBSCAN analysis) | Absolute PC loadings for al principal components for each feature represented as a heatmap. (.png) | FO and IO |
| **Feature_importance_TP0.xlsx, Feature_importance_TP1.xlsx, Feature_importance_TP2.xlsx, …** | PCA (for k-means and DBSCAN analysis) | PC loadings for al principal components for each feature represented as a matrix in an .xlsx file | FO and IO |
| **Feature_importance_TP0.png, Feature_importance_TP1.png, Feature_importance_TP2.png, …** | PCA (for k-means and DBSCAN analysis) | PC loadings for al principal components for each feature represented as a heatmap. (.png) | FO and IO |
| **timepoint_0.xlsx, timepoint_1.xlsx, timepoint_2.xlsx, …** | PCA (for k-means and DBSCAN analysis) | Multivariate tables per timepoint labelled with condition name. Rows represent individual spheroids and columns represent different features (.xlsx) | FO and IO |
| **CLUSTER_timepoint_0.xlsx, CLUSTER_timepoint_1.xlsx, CLUSTER_timepoint_2.xlsx, …** | PCA (for k-means and DBSCAN analysis) | Multivariate tables per timepoint labelled with cluster ID. Rows represent individual spheroids and columns represent different features (.xlsx) These are also generated for the secondary PCA2. for these, file names have "NEW" as prefix | FO and IO |
| **MINMAX_timepoint_0.xlsx, MINMAX_timepoint_1.xlsx, MINMAX_timepoint_2.xlsx, …** | PCA (for k-means and DBSCAN analysis) | Min-max scaled multivariate tables per timepoint labelled with condition name. Rows represent individual spheroids and columns represent different features. (.xlsx) | FO and IO |
| **Kmeans_inertia_TP0.png, Kmeans_inertia_TP1.png, Kmeans_inertia_TP2.png, ...** | PCA (for k-means analysis) | K-means inertia plot displaying how the optimal number of clusters can be estimated by the elbow method. (.png) These plots are also generated for the secondary PCA2. for these, plot names have "NEW" as prefix | FO |

| | | | |
|---|---|---|---|
| **Kmeans_CH_TP0.png, Kmeans_CH_TP1.png, Kmeans_CH_TP2.png, ...** | PCA (for k-means analysis) | Calinski–Harabasz index across increasing cluster numbers, used to identify the optimal k-means solution. These plots are also generated for the secondary PCA2. for these, plot names have "NEW" as prefix | FO |
| **PC1and2_Abs_Feature_imp_TP0.png, PC1and2_Abs_Feature_imp_TP1.png, PC1and2_Abs_Feature_imp_TP2.png, ...** | PCA (for k-means analysis) | Absolute feature-importance scores for PC1 and PC2 at every timepoint, showing the contribution of each feature to the principal components driving phenotypic variation. (png, heatmap) | FO |
| **subpl_Abs_Feature_imp_TP0.png, subpl_Abs_Feature_imp_TP1.png, subpl_Abs_Feature_imp_TP2.png, ...** | PCA (for k-means analysis) | Similar as PC1and2_Abs_Feature_imp_TPX.png. In this plot, the mean of PC1 and PC2 is added enable illustrative comparison with the weighted mean. (.png, heatmap) | FO |
| **wMean_all PC_TP0.png, wMean_all PC_TP1.png, wMean_all PC_TP2.png, ...** | PCA (for k-means and DBSCAN analysis) | Heatmap (.png) of the Feature Importance for each feature at every timepoint. The FI is the weighted mean of the absolute loadings of all PCs. | FO |
| **PCA_var TP0.png, PCA_var TP1.png, PCA_var TP2.png, ...** | PCA (for k-means and DBSCAN analysis) | Bar plots displaying the explained variance per principal component at every timepoint. (.png) The explained variance shown is the weight used in the feature importance calculation. | FO |
| **PCA_COND_timepoint_0.png, PCA_COND_timepoint_1.png, PCA_COND_timepoint_2.png, ...** | PCA (for k-means and DBSCAN analysis) | PCA plots (.png) showing PC1 and PC2. Individual points are spheroids coloured by condition. These plots are also generated for the secondary PCA2. for these, plot names have "NEW" as prefix | FO |
| **Kmeans_Clustering_ConvexHull_TP0.png, Kmeans_Clustering_ConvexHull_TP1.png, Kmeans_Clustering_ConvexHull_TP2.png, ...** | PCA (for k-means analysis) | PCA plots (.png) showing PC1 and PC2. Individual points are spheroids coloured by condition. PCA plots are overlayed with k-means cluster convex hulls, visually showing the identified clusters. These plots are also generated for the secondary PCA2. for these, plot names have "NEW" as prefix | FO |

| | | | |
|---|---|---|---|
| **DBSCAN_Clustering_timepoint_0.png, DBSCAN_Clustering_timepoint_1.png, DBSCAN_Clustering_timepoint_2.png, ...** | PCA (for DBSCAN analysis) | PCA plots (.png) showing PC1 and PC2. Individual points are spheroids coloured by DBSCAN cluster. These plots are also generated for the secondary PCA2. for these, plot names have "NEW" as prefix | FO |
| **K-distance_Plot_timepoint_0.png, K-distance_Plot_timepoint_1.png, K-distance_Plot_timepoint_2.png, …** | PCA (for DBSCAN analysis) | K-distance plot at timepoint 0, used to estimate the optimal epsilon (ε) parameter for DBSCAN clustering. These plots are also generated for the secondary PCA2. for these, plot names have "NEW" as prefix | FO and IO |
| **PCA\ClusterBoxplots** | PCA | SF | SF |
| **Area_Single_Cells_boxplot_TP0.png, Area_Single_Cells_boxplot_TP1.png, Area_Single_Cells_boxplot_TP2.png, Area_Single_Cells_boxplot_TP3.png, Area_Single_Cells_boxplot_TP4.png, …** | PCA\ ClusterBoxplots (for k-means and DBSCAN analysis) | Boxplots (.png) of each feature for every timepoint stratified by cluster ID | FO |
| **Plots\BAR** | Plots | SF | SF |
| **Plots\BOX** | Plots | SF | SF |
| **Plots\Heatmap** | Plots | SF | SF |
| **Plots\SCAT** | Plots | SF | SF |
| **areaSingleCells t1 Boxplot with CLD.png, areaSingleCells t2 Boxplot with CLD.png, areaSingleCells t3 Boxplot with CLD.png, areaSingleCells t4 Boxplot with CLD.png, …** | Plots\BOX | Boxplots (.png) of each feature for every timepoint except t0. On this plot, statistical results are shown as compact letter display. | FO |
| **areaSingleCells t1 NS network.png, areaSingleCells t2 NS network.png, areaSingleCells t3 NS network.png, areaSingleCells t4 NS network.png, …** | Plots\BOX | Non-significance networks (.png) for each feature at each timepoint, illustrating the statistical relationships between conditions. | FO |
| **Dunn--Heatmap_aspectRatio t3.png, Dunn--Heatmap_aspectRatio t4.png, Dunn--Heatmap_areaSingleCells t1.png, Dunn--Heatmap_areaSingleCells t2.png, Dunn--Heatmap_areaSingleCells t3.png, …** | Plots\Heatmap | Heatmap plots (.png), visualizing the p-value matrix (incl. *, **, ***, ****, ns) of Stats\Dunn and Stats\Tukey results for all features at every timepoint except t0. | FO |

| | | | |
|---|---|---|---|
| **Barplot Protrusion_count_Protr. count.png, Barplot Cell_count_Cell count.png, Barplot Invasion_Depth_Distance.png, Barplot Invasion_Depth_Invasion depth.png, Barplot Invasion_Depth_Invasion Ratio.png, Barplot Invasion_Mode_Index_IMI.png, …** | Plots\BAR | Contains combined bar plots (.png) for every feature. The plot consists of all subsequent timepoints for all conditions per feature. This makes it easy to assess increase of decrease of a feature over time. | FO |
| **Scatterplot Invasion_Mode_Index_IMI.png, Scatterplot Protrusion_count_Protr. count.png, Scatterplot Cell_count_Cell count.png, Scatterplot Invasion_Depth_Distance.png, Scatterplot Invasion_Depth_Invasion depth.png, Scatterplot Invasion_Depth_Invasion Ratio.png, …** | Plots\SCAT | Similar as the bar plots. Contains combined bar plots (.png) for every feature. The plot consists of all subsequent timepoints for all conditions per feature with individual spheroid values overlayed as scatter to visualize the value distribution. This makes it easy to assess increase of decrease of a feature over time. | FO |
| **Anova_KruskalWallis.txt** | Stats | This file contains A) which parameters do and or do not have a normal distribution according to the results in Stats\Normality. B) H-statistic and p-value of the ANNOVA/Kruskal-Wallis analysis for each feature for each timepoint | FO |
| **Stats\Dunn** | Stats | SF | SF |
| **Stats\Normality** | Stats | SF | SF |
| **Stats\QQ** | Stats | SF | SF |
| **Stats\Tukey** | Stats | SF | SF |
| **Cell_count t1.xlsx, aspectRatio t3.xlsx, aspectRatio t4.xlsx, areaSingleCells t1.xlsx, areaSingleCells t2.xlsx, areaSingleCells t3.xlsx, …** | Stats\Dunn | .xlsx files containing a p-value matrix for all features at all timepoints excluding t0 that were analysed using Dunn post-hoc test for multiple comparisons after Kruskal-Wallis (non-parametric, non-normal distribution) | FO and IO |
| **areaSingleCells t4.xlsx, areaSpheroid t1.xlsx, areaSpheroid t2.xlsx, areaSpheroid t3.xlsx, areaSpheroid t4.xlsx, areaTotal t1.xlsx, …** | Stats\Tukey | .xlsx files containing a p-value matrix for all features at all timepoints excluding t0 that were analysed using Tukey post-hoc test for multiple comparisons after one-way ANOVA (parametric, normal distribution) | FO and IO |

| | | | |
|---|---|---|---|
| **Normality.txt** | Stats\Normality | This file contains the results of the Shapiro-Wilk Test including W, p-value and Normality (T/F) for each condition for each feature at every timepoint. A feature is considered as normally distributed if all individual tested conditions have a normal distribution | FO and IO |
| **Overall Normality.xlsx** | Stats\Normality | This file contains the results the overall normality assessment of each feature base on the Shapiro-Wilk test results of each individual condition. This is a Boolean list for each feature at every timepoint. True = Normal distribution, False = Non-normal distribution | FO and IO |
| **QQ-plot areaSingleCells t1.csv.png, QQ-plot areaSingleCells t2.csv.png, QQ-plot areaSingleCells t3.csv.png, QQ-plot areaSingleCells t4.csv.png, QQ-plot areaSpheroid t0.csv.png, QQ-plot areaSingleCells t0.csv.png, …** | Stats\QQ | Contains QQ plots (.png) of all features for every timepoint. Each plot contains a separate subplot for each condition. This is an optional step to visually represent normality in each condition, however generating this takes time and storage. | FO |

***Table S4. SImBA segmentation and performance metrics***

Comprehensive overview of datasets used to assess SImBA segmentation performance across different microscope platforms, imaging modalities, and cell lines. For each dataset, the microscope used for image acquisition, data source, cell line, imaging mode, SImBA segmentation method, Gaussian sigma/saturated pixels, total number of analysed images, number of correctly segmented spheroids (CS) by visual inspection of the segmentation, and percentage of correct segmentation are listed. Growth and invasion assays, as well as phase-contrast, brightfield, and fluorescence datasets, are included. The table illustrates the broad applicability and high segmentation accuracy of SImBA across heterogeneous imaging conditions. BF, brightfield; PH, phase contrast; FL, fluorescence; NAT, no auto-threshold; DB, double blur; SP: Saturated pixels (only for FL), TNAI: total number of analysed spheroids, CS, correctly segmented spheroids.

| Microscope | Source | Cell line | Imaging mode | SImBA Segmentation Method | Sigma/ SP | TNA S | CS | % CS |
|---|---|---|---|---|---|---|---|---|
| ***Incucyte ZOOM*** | Laboratory of Experimental Cancer Research (2025). IncucyteZOOM.zip. figshare. Dataset. https://doi.org/10.6084/m9.figshare.27438526.v1 | 4T1 | PH | GROWTH; Double Blur (DB) | 5.5 | 742 | 741 | 99.87 |
| | | A549 | PH | GROWTH; Default (NAT) | 6.5 | | | |
| | | D2A1 | PH | GROWTH; Default (NAT) | 4.5 | | | |
| | | HCT116 | PH | GROWTH; Default (NAT) | 4.5 | | | |
| | | HepG2 | PH | GROWTH; Double Blur (DB) | 5.5 | | | |
| | | PANC1 | PH | GROWTH; Default (NAT) | 4.5 | | | |
| | | SKOV3 | PH | GROWTH; Default (NAT) | 5.5 | | | |
| ***Axiovert 200 M*** | Laboratory of Experimental Cancer Research (2025). Axiovert200M.zip. figshare. Dataset. https://doi.org/10.6084/m9.figshare.27393225.v1 | BT474 | BF | GROWTH; Default (NAT) | 4.5 | 429 | 429 | 100 |
| | | CAL33 | BF | GROWTH; Double Blur (DB) | 8.5 | | | |
| | | FaDu | BF | GROWTH; Double Blur (DB) | 8.5 | | | |
| | | HEK293 | BF | GROWTH; Default (NAT) | 4.5 | | | |
| | | HOP62 | BF | GROWTH; Default (NAT) | 6.5 | | | |
| | | HSC4 | BF | GROWTH; Double Blur (DB) | 7.5 | | | |
| | | MDA-MB-231 | BF | GROWTH; Default (NAT) | 4.5 | | | |
| | | NCIH226 | BF | GROWTH; Default (NAT) | 5.5 | | | |

| | | | | | | | | |
|---|---|---|---|---|---|---|---|---|
| | | NCIH460 | BF | GROWTH; Default (NAT) | 4.5 | | | |
| | | OVCAR8 | BF | GROWTH; Default (NAT) | 4.5 | | | |
| | | U138M6 | BF | GROWTH; Default (NAT) | 4.5 | | | |
| | | U251M6 | BF | GROWTH; Default (NAT) | 4.5 | | | |
| ***Cytation 5*** | Laboratory of Experimental Cancer Research (2025). Cytation5.zip. figshare. Dataset. https://doi.org/10.6084/m9.figshare.27393219.v1 | CT5.3hTERT | BF | GROWTH; Default (NAT) | 4.5 | 66 | 66 | 100 |
| ***Opera Phenix*** | Laboratory of Experimental Cancer Research (2025). OperaPhenix.zip. figshare. Dataset. https://doi.org/10.6084/m9.figshare.27390402.v1 | Huh7 | BF | GROWTH; Default (NAT) | 4.5 | 30 | 30 | 100 |
| | | LNCAP | BF | GROWTH; Default (NAT) | 4.5 | | | |
| ***Olympus IX05*** | Laboratory of Experimental Cancer Research (2025). OlympusIX05.zip. figshare. Dataset. https://doi.org/10.6084/m9.figshare.27390411.v1 | A549-MRC5 | BF | GROWTH; Double Blur (DB) | 6.5 | 88 | 80 | 90.91 |
| | | H1299 | BF | GROWTH; Double Blur (DB) | 5.5 | | | |
| | | MRC5 | BF | GROWTH; Double Blur (DB) | 6.5 | | | |
| | | VCAP | BF | GROWTH; Double Blur (DB) | 5.5 | | | |
| ***Olympus IX81*** | This study | Hep3B | PH | INVASION; Default (NAT) | 6.5 | 1970 | 1912 | 97.06 |
| | | SNU423 | PH | INVASION; Default (NAT) | 6.5 | | | |
| | | Huh7 | PH | INVASION; Default (NAT) | 6.5 | | | |
| | | Hela | PH | INVASION; Default (NAT) | 6.5 | | | |
| | | Hepa1-6 | PH | INVASION; Default (NAT) | 6.5 | | | |
| | | MDA-MD-231 | PH | INVASION; Default (NAT) | 6.5 | | | |
| | | HT1080 | PH | INVASION; Default (NAT) | 6.5 | | | |
| | | SK_MEL_28 | PH | INVASION; Default (NAT) | 6.5 | | | |
| ***Olympus*** | This study | Hep3B | FL | INVASION; Fluorescence | 5.5 | 171 | 170 | 99.42 |

| | | | | | | | | |
|---|---|---|---|---|---|---|---|---|
| ***IX81*** | | SNU423 | FL | INVASION; Fluorescence | 5.5 | | | |
| | | | | | **TOTAL** | **3496** | **3428** | **98.05** |

*Table S5. Overview of metadata for case study 1 and 2*

Overview of case study metadata. The table lists, for each condition, the case study, cell line, RRID Cellosaurus identifier, spheroid formation time before embedding/imaging, treatment condition, total imaging duration, interval and number of time points, toxicity staining, number of analysed spheroids and the total of analysed images. Case study 1 compares invasion phenotypes across eight spheroid-forming cell lines of diverse origin under untreated conditions. Case study 2 examines dose-dependent sorafenib (SFN) responses in Hep3B and SNU-423 spheroids. In total, 460 spheroids were included across both case studies. CS: case study, $T_f$: final timepoint, NTP: number of timepoints, NS: number of spheroids, NIC: number of images per condition

| CS | Cell line | RRID Cellosaurus | Spheroid formation time (h) | Treatment | $T_f$ (h) | Time interval (h) | NTP | Cytotoxicity staining | NS | NIC |
|---|---|---|---|---|---|---|---|---|---|---|
| **CS 1** | *Hep3B* | CVCL_0326 | 24 | None | 48 | 24 | 3 | NO | 30 | 90 |
| **CS 1** | *Huh-7* | CVCL_0336 | 24 | None | 48 | 24 | 3 | NO | 32 | 96 |
| **CS 1** | *SNU-423* | CVCL_0366 | 24 | None | 48 | 24 | 3 | NO | 37 | 111 |
| **CS 1** | *Hela* | CVCL_0030 | 24 | None | 48 | 24 | 3 | NO | 48 | 144 |
| **CS 1** | *Hepa 1-6* | CVCL_0327 | 24 | None | 48 | 24 | 3 | NO | 20 | 60 |
| **CS 1** | *MDA-MB-231* | CVCL_0062 | 72 | None | 48 | 24 | 3 | NO | 29 | 87 |
| **CS 1** | *HT-1080* | CVCL_0317 | 72 | None | 48 | 24 | 3 | NO | 37 | 111 |
| **CS 1** | *SK-MEL-28* | CVCL_0526 | 72 | None | 48 | 24 | 3 | NO | 34 | 102 |
| **CS 2** | *Hep3B* | CVCL_0326 | 24 | SFN 0µM | 96 | 24 | 5 | YES | 25 | 95 |
| **CS 2** | *Hep3B* | CVCL_0326 | 24 | SFN 10µM | 96 | 24 | 5 | YES | 19 | 85 |
| **CS 2** | *Hep3B* | CVCL_0326 | 24 | SFN 15µM | 96 | 24 | 5 | YES | 17 | 55 |
| **CS 2** | *Hep3B* | CVCL_0326 | 24 | SFN 25µM | 96 | 24 | 5 | YES | 11 | 75 |
| **CS 2** | *Hep3B* | CVCL_0326 | 24 | SFN 50µM | 96 | 24 | 5 | YES | 15 | 70 |
| **CS 2** | *SNU-423* | CVCL_0366 | 24 | SFN 0µM | 96 | 24 | 5 | YES | 14 | 75 |
| **CS 2** | *SNU-423* | CVCL_0366 | 24 | SFN 5µM | 96 | 24 | 5 | YES | 15 | 105 |
| **CS 2** | *SNU-423* | CVCL_0366 | 24 | SFN 10µM | 96 | 24 | 5 | YES | 21 | 85 |
| **CS 2** | *SNU-423* | CVCL_0366 | 24 | SFN 15µM | 96 | 24 | 5 | YES | 17 | 95 |
| **CS 2** | *SNU-423* | CVCL_0366 | 24 | SFN 25µM | 96 | 24 | 5 | YES | 19 | 100 |
| **CS 2** | *SNU-423* | CVCL_0366 | 24 | SFN 50µM | 96 | 24 | 5 | YES | 20 | 95 |
| | | | | | | | | **TOTAL** | **460** | **1736** |
